# The P450 Sterol Side Chain Cleaving Enzyme (P450_scc_) for Digoxin Biosynthesis in the Foxglove Plant Belongs to the CYP87A Family

**DOI:** 10.1101/2022.12.29.522212

**Authors:** Emily Carroll, Baradwaj Ravi Gopal, Indu Raghavan, Zhen Q. Wang

## Abstract

Digoxin from the foxglove plant is a commonly prescribed plant natural product for treating heart failure and atrial fibrillation. Despite its medicinal prominence, how foxglove synthesizes digoxin is largely unknown, especially the cytochrome P450 sterol side chain cleaving enzyme (P450_scc_), which catalyzes the first and rate-limiting step in this pathway. Here we report the identification of the foxglove P450_scc_, the gatekeeping enzyme that channels sterols to digoxin. This enzyme converts both cholesterol and campesterol to pregnenolone, thus explaining how pregnenolone is synthesized in plants. Phylogenetic analysis indicates that this enzyme arose from a duplicated *CYP87A* gene and does not share clear homology with the mammalian P450_scc_. Identifying this long-speculated plant P450_scc_ enzyme suggests that the digoxin biosynthetic pathway starts from both cholesterol and phytosterols, instead of just cholesterol as previously thought. The identification of this gatekeeping enzyme is a key step towards complete elucidation of digoxin biosynthesis and expanding the therapeutic applications of digoxin analogs in future work.

## Introduction

Cardiac glycosides extracted from the foxglove plant *Digitalis lanata* have been used for treating congestive heart failure since 1785^1^. One cardiac glycoside, digoxin, is widely prescribed for treating heart failure and atrial fibrillation and is listed by the World Health Organization as an essential medicine^2^. There were about 400,000 patients prescribed digoxin in the United States in 2020, making it one of the most prescribed plant natural products^3^. Recent research has broadened the medicinal application of digoxin and other cardiac glycosides for treating viral infection, inflammation, cancer (Anvirzel, Phase I clinical trial), hypertension (Rostafuroxin, Phase II clinical trial), and neurodegenerative diseases^4–11^.

One aspect of digoxin and other cardiac glycosides that limits their broader use is their toxicity^10,11^. Digoxin has a narrow therapeutic range of 0.8-2.0 ng/mL and exceeding this range leads to severe toxicity^12,13^. Modifications of functional groups in the digoxin molecule change the binding affinity of digoxin to its receptor, the Na^+^/K^+^ ATPase pump, and improve its pharmacokinetic profile^7,14^. However, these structural modifications are difficult to achieve as digoxin is extracted from the leaves of *D. lanata* at only 0.06% of dry weight^15^. Total synthesis of cardiac glycosides is challenging due to their structural complexity^16^. Alternatively, synthesizing digoxin in engineered microbes would potentially increase its yield and facilitate structural modification. Such an endeavor requires transforming the microbial host with all genes responsible for synthesizing cardiac glycosides. However, most of these genes are unknown.

Due to the prominence of digoxin in medicine, the study of the cardiac glycoside biosynthetic pathways dates back to the 1960s. Radiolabeling studies suggested cholesterol to be the precursor for digoxin^17^. While this is generally accepted, controversies remain since cholesterol is only a minor sterol in plants. The exact biosynthetic pathway of digoxin remains enigmatic more than half a century after the initial ratio-labeling works^18^. The hypothetical cardiac glycoside biosynthetic pathway starts with cholesterol which undergoes nine enzyme-catalyzed steps to digoxigenin, the steroidal scaffold of digoxin^18^. Currently, the only known enzymes in the pathway are 3*β*-hydroxysteroid dehydrogenase (3*β*HSD) and progesterone-5*β-*reductase (P5*β*R and P5*2β*R)^19–21^. The first and rate-limiting enzyme, cytochrome P450 sterol side chain leaving enzyme (P450_scc_), along with all of the other enzymes have not been identified yet^18,22^. The foxglove P450_scc_ is thought to convert cholesterol to pregnenolone through a reaction identical to the mammalian P450_scc_, which catalyzes the rate-limiting step in animal steroid hormone synthesis^23^. However, this critical enzyme has not been isolated and characterized since its first description by Pilgrim in 1972^24^. Hence, the direct sterol precursor for cardiac glycosides biosynthesis remains ambiguous as indirect evidence suggests that phytosterols, main sterols in plants, may also be precursors for cardiac glycosides^17,24–28^.

In this study we present a high-quality transcriptome of *D. lanata* which aided the identification of the foxglove P450_scc_. Characterizing the foxglove P450scc by heterologous expression in tobacco and yeast validated its sterol cleaving activity and substrate preference, and reveals the identity of sterol precursors for digoxin biosynthesis. Phylogenetic analysis suggests possible evolutionary mechanism of this enzyme. The foxglove P450_scc_ identified here is the first P450_scc_ reported in plants and does not share clear homology with the animal P450_scc_.

## Results

### Transcriptome assembly and annotation

RNA from six leaf and six root samples were pooled together to generate a high-quality reference transcriptome of *D. lanata*. Since neither the genome nor transcriptome is available, we performed the *de novo* assembly of the transcriptome from 173,448,870 Illumina raw reads with an average length of ∼100 bp. The assembled transcriptome contains 317,983 transcripts with an N50 of 1,712 bp (Table 1, Supplementary Figure S1). The Benchmarking Universal Single-Copy Orthologs (BUSCO) score for the transcriptome was 94.6% and the raw reads mapped to the transcriptome assembly showed 99.36% alignment indicating that the transcriptome was near complete. 190,755 transcripts with at least 300-bp long were annotated using publicly available databases including the NCBI non-redundant protein database (nr) and the UniProt database, each annotated 84.6% and 48.3% of the 190,755 transcripts, respectively^29,30^. The transcripts were found to match mostly with genes of other Lamiales species, including *Sesamum indicum*, which covered 75.3% of the transcriptome, as expected (Supplementary Figure S1). 113,221 non-redundant unigenes were also categorized by Gene Ontology (GO) and Kyoto Encyclopedia of Genes and Genomes (KEGG) pathway classification^31,32^ (Supplementary Figure S2). KEGG analysis revealed 5,683 unigenes in 412 KEGG pathways, among which were pathways for terpenoid and steroid biosynthesis (Supplementary Figure S3). UniProt annotation identified 7,517 transcription factors and regulators, 4,226 protein kinases, and 22,549 simple sequence repeats (SSR) as genetic markers (Table 1, Supplementary Figure S4, Supplementary Table S1). The annotated transcriptome presented here provides a comprehensive representation of transcripts within the root and leaf tissues of *D. lanata*.

**Table 1.**
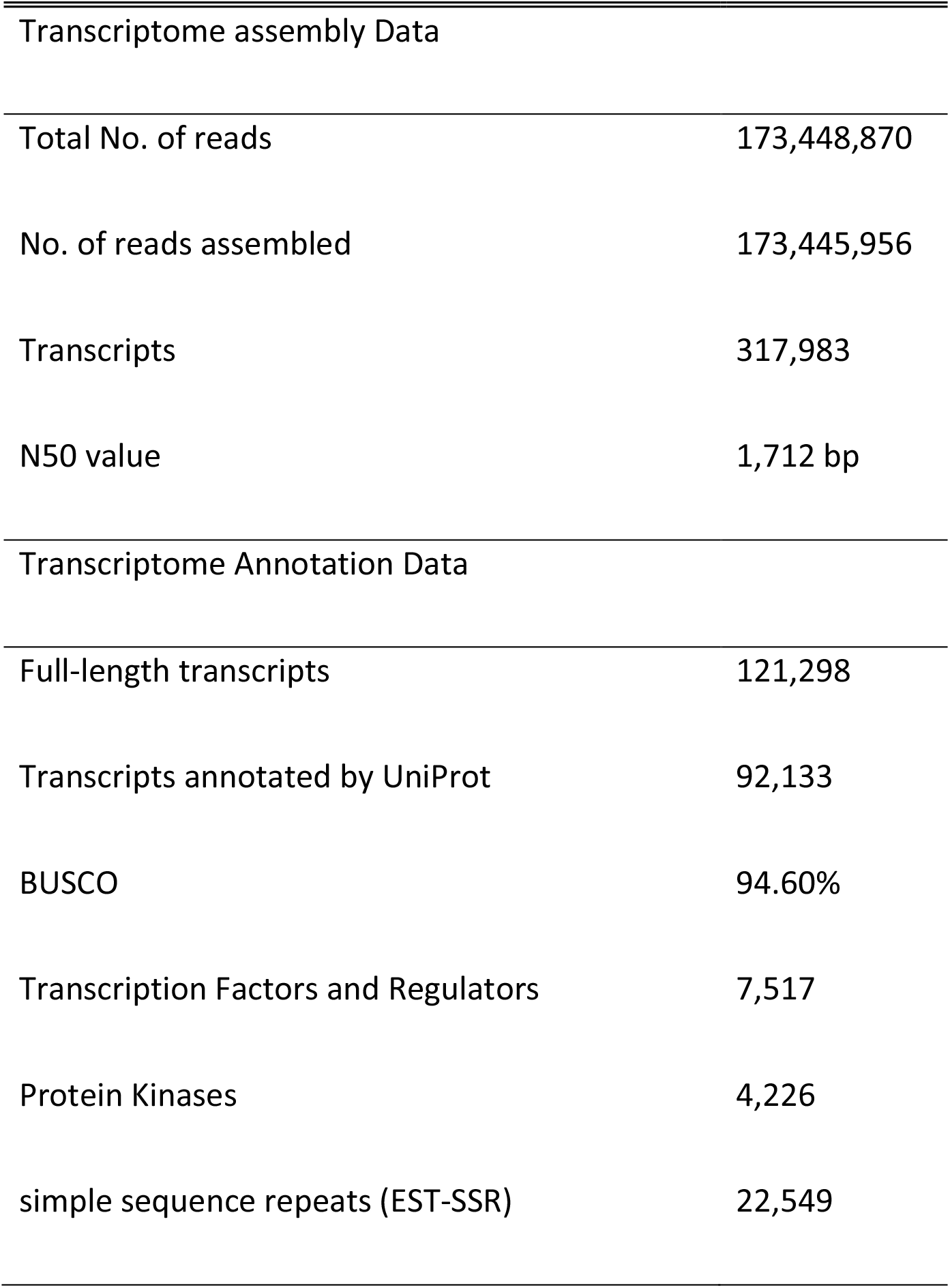
Summary of Transcriptome data

|  |  |
| --- | --- |
| Transcriptome assembly Data |  |
| Total No. of reads | 173,448,870 |
| No. of reads assembled | 173,445,956 |
| Transcripts | 317,983 |
| N50 value | 1,712 bp |
| Transcriptome Annotation Data |  |
| Full-length transcripts | 121,298 |
| Transcripts annotated by UniProt | 92,133 |
| BUSCO | 94.60% |
| Transcription Factors and Regulators | 7,517 |
| Protein Kinases | 4,226 |
| simple sequence repeats (EST-SSR) | 22,549 |

### Genes for sterol biosynthesis are differentially expressed in leaves and roots

Digoxin biosynthesis is widely considered to begin with cholesterol but indirect evidence suggests phytosterols may also be substrates^18,24^. Phytosterols including campesterol, stigmasterol, and sitosterol are 24-alkyl sterols unique to plants^33^. Inhibiting the 24-methyltransferase in the phytosterol pathway increased cholesterol levels but decreased the amount of cardiac glycosides in foxglove plants^27^. Our RNA-seq analysis identified key genes of the cholesterol and phytosterol pathway that are differentially expressed in the leaves and roots of *D. lanata*.

Since cardiac glycosides are only present in leaves but not roots (Figure 1A), we asked if the genes for phytosterol and cholesterol biosynthesis are overexpressed in leaves. Indeed, genes encoding rate-limiting enzymes in phytosterol and cholesterol pathways are overexpressed in leaves (Figure 1B). Squalene epoxidase (SQE), a rate-limiting step in sterol biosynthesis, is induced in leaves^34^. Sterol side-chain reductase (SSR1) catalyzes the first step in the cholesterol pathway and the last step in the campesterol/*β*-sitosterol pathway in leaves^35^. SSR1 is a known bottleneck enzyme^36^ in plant sterol biosynthesis and is also overexpressed in *D. lanata* leaves. SMO3 duplicated from SMO1 and unique to the cholesterol pathway is also induced in leaves. It catalyzes the rate-limiting step of 4-methyl elimination in the cholesterol pathway^28^. Indeed, *D. lanata* leaves have higher levels of cholesterol than roots even though the total sterols in these two tissues are comparable (Supplementary Figure S5). Another overexpressed enzyme is the sterol C-14 reductase (C14-R)^37^, a shared enzyme between phytosterol and cholesterol pathways. The transcriptome analysis indicates that key enzymes in the cholesterol and phytosterol pathways are overexpressed in leaves.

**Figure 1:**
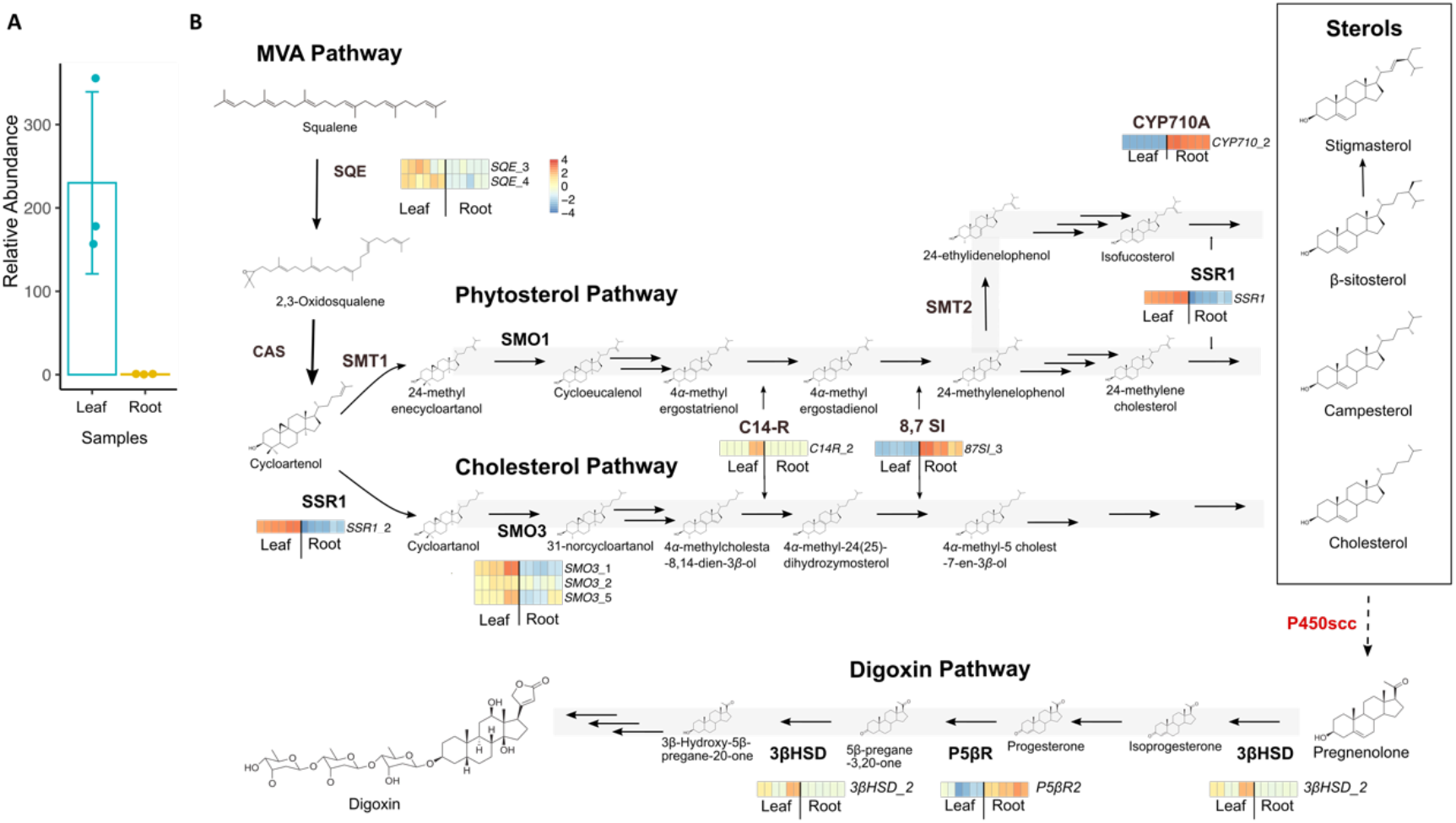
Differential expression analysis of genes in the sterol and digoxin biosynthetic pathways in *D. lanata* leaf and root samples. A. Total lanatosides including lanatoside A, B, C, and E in leaf and root tissues of *D. lanata* seedlings relative to digoxin-d3 internal standard. The data are normalized by dry weight and represent average ± SD of three biological replicates. B. Sterol and digoxin biosynthesis pathway genes that are differentially expressed in *D. lanata* leaf and root tissues. Six replicates of leaf and root samples are shown in the heat maps. Genes that do not have corresponding heat maps shown are not differentially expressed. SQE: squalene epoxidase; CAS: cycloartenol synthase; SMO: C-4 sterol methyl oxidase, SMT: sterol C-24 methyl transferase; SSR 1: sterol side-chain reductase 1. C14-R: sterol C-14 reductase; 8,7 SI: sterol 8,7 isomerase; P450_scc_: cytochrome P450 sterol side-chain cleaving; 3*β*HSD: 3*β*-hydroxysteroid dehydrogenase; P5*β*R: progesterone-5*β*-reductase.

Although most enzymes in the digoxin biosynthetic pathway are unknown, analysis of the three known enzymes shows that only 3*β*HSD was overexpressed in *D. lanata* leaves. While P5*β*R is constitutively expressed in both tissues, P5*β*R2 is more abundant in roots, consistent with the previous reports that P5*β*R2 is expressed at low levels in leaves in the absence of stress^21^. Since digoxin and sterols are triterpene derivatives, we also analyzed the differential expression of terpenoid biosynthetic genes (Supplementary Figure S6). The methylerythritol phosphate (MEP) pathway for terpenoid synthesis and triterpene pathways are induced in leaves, agreeing with the compartmentalization of the MEP pathway in the chloroplasts^38^.

### Identification of two candidate *D. lanata* P450_scc_

The first step of the digoxin pathway involves the cleavage of a sterol by a P450_scc_ enzyme to produce pregnenolone^18^. Interpro scan identified 438 enzymes that were annotated within the cytochrome P450 (CYP) Pfam family (PF00067)^39,40^. CYPs from *Arabidopsis thaliana* and CYPs found in *D. lanata* transcriptome were used to construct a phylogenetic tree for CYP subfamily classification (Supplementary Figure S7). Differential expression analysis identified 104 CYP transcripts that were overexpressed in the leaves (Supplementary Figure S8). Amongst these CYPs only those that were members of subfamilies relevant to sterol/brassinosteroid biosynthesis were included in future analysis. Thirteen of such full-length CYP transcripts were identified as potential *P450*_*scc*_ (Figure 2A). We focused on *DlCYP87A4* and *DlCYP90A1* because *DlCYP87A4* was highly induced in leaves and CYP90A1 is known to oxidize the C3 group of 22*(S)*-hydroxycampesterol^41^. Another CYP87A subfamily member, *DlCYP87A3*, was also highly induced in the leaf and shares a 97.4% protein identity to *DlCYP87A4* but could not be amplified from cDNA. qRT-PCR confirmed that *DlCYP87A4* was expressed much higher in leaves compared to *DlCYP90A1* (Fig 2B). These two transcripts were identified as *P450*_*scc*_ candidates and cloned from cDNA for functional validation by tobacco transient expression assay.

**Figure 2:**
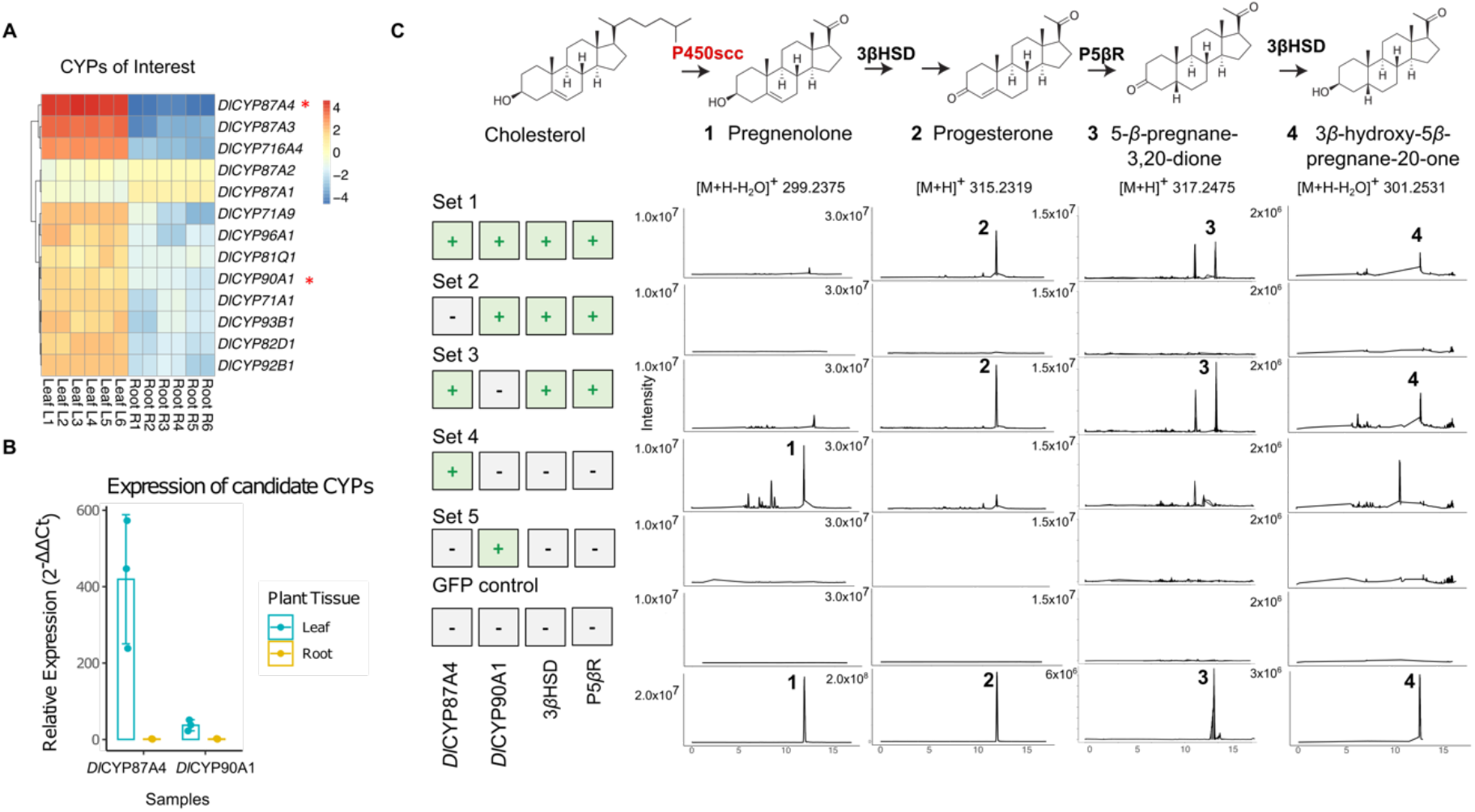
Identification and characterization of candidate *D. lanata* P450_scc_ in tobacco. A. Differential expression analysis of candidate P450_scc_ in root and leaf tissues of *D. lanata*. Six replicates of leaf and root samples are shown in the heat maps. B. qRT-PCR quantified relative expressions of top candidate genes, *Dl*CYP87A4 and *Dl*CYP90A1, in root and leaf samples. C. LC/MS data from tobacco leaves transiently expressing candidate genes in various combinations. Set 1 contained *Dl*CYP87A4, *Dl*CYP90A1, 3*β*HSD and P5*β*R. Set 2 contained *Dl*CYP90A1, 3*β*HSD, and P5*β*R. Set 3 contained *Dl*CYP87A4, 3*β*HSD and P5*β*R. Set 4 contained *Dl*CYP87A4 only. Set 5 contained *Dl*CYP90A1 only. GFP control: negative control expressing a green fluorescent protein (GFP) in tobacco. Authentic standards of the four expected pathway intermediates are shown in the bottom panel.

### Tobacco expression identifies *D. lanata* P450_scc_ as CYP87A4

In order to test the two candidate enzymes, we employed the tobacco transient expression experiment. Tobacco does not produce digoxin or any of the pathway intermediates, but it has sterol substrates for the P450_scc_^37.^ Therefore, it is a good system for functionally characterizing the P450_scc_ enzymes. The two candidates, *Dl*CYP87A4 and *Dl*CYP90A1, along with the two known pathway enzymes, 3*β*HSD and P5*β*R, were all expressed together in tobacco leaves. Following their expression, products of these enzymes including progesterone (compound 2), 5*β-*pregnane-3,20-one (compound 3), and 3*β*-hydroxy-5*β*-pregnane-20-one (compound 4), were detected (Figure 2C, set 1, Supplementary Figure S9). Omitting the *Dl*CYP87A4 abolishes the reactions (set 2) whereas taking out the *Dl*CYP90A1 (set 3) had no effect. The direct product of P450_scc_, pregnenolone (compound 1), was not detected potentially due to its quick turnover through 3*β*HSD. In fact, *D. lanata* leaves do not produce detectable amounts of pregnenolone but do produce the downstream pathway intermediates (Supplementary Figure S10). Expressing *Dl*CYP87A4 alone resulted in the production of pregnenolone (set 4) whereas expressing *Dl*CYP90A1 alone (set 5) did not produce any of this product. These data strongly support the hypothesis that *Dl*CYP87A4 is a functional P450_scc_ of the digoxin pathway that cleaves a plant sterol to produce pregnenolone.

### Determining the sterol substrates of CYP87A4

The tobacco expression system does not allow the determination of sterol substrates of *D. lanata* P450_scc_ since tobacco contains a mixture of cholesterol and phytosterols. In order to understand the substrate specificity of *D. lanata* CYP87A4, we expressed this enzyme along with an *Arabidopsis* cytochrome P450 reductase 2 (ATR2) as the redox partner^42^ in *S. cerevisiae*. Since feeding yeast with different sterols is challenging due to their hydrophobicity, we turned to the previously engineered yeast strains that produces various sterols, including cholesterol, campesterol, 7-dehydrocholesterol, and desmosterol, as major sterols in their cell membranes (Supplementary Figure S11)^37^. The wildtype yeast has ergosterol as its main sterol. These sterols differ slightly in their side-chain and B-ring structures (Figure 3A). When expressing the D. *lanata* CYP87A4 and ATR2, only the yeast strains containing campesterol and cholesterol respectively produced pregnenolone (Figure 3B, Supplementary Figure S12). Pregnenolone is toxic to yeast and as a result yeast acetylates pregnenolone to detoxify it, generating pregnenolone acetate (compound 5) as a byproduct^43^. We also expressed the human P450_scc_ (CYP11A1) with its redox partners in yeast as a positive control and only the cholesterol-producing yeast generated pregnenolone, as expected (Figure 3B)^23^. These data indicate that both campesterol and cholesterol are precursors of the digoxin pathway and highlights the promiscuous nature of the *Dl*CYP87A4 now identified as the *D. lanata* P450_scc_.

**Figure 3:**
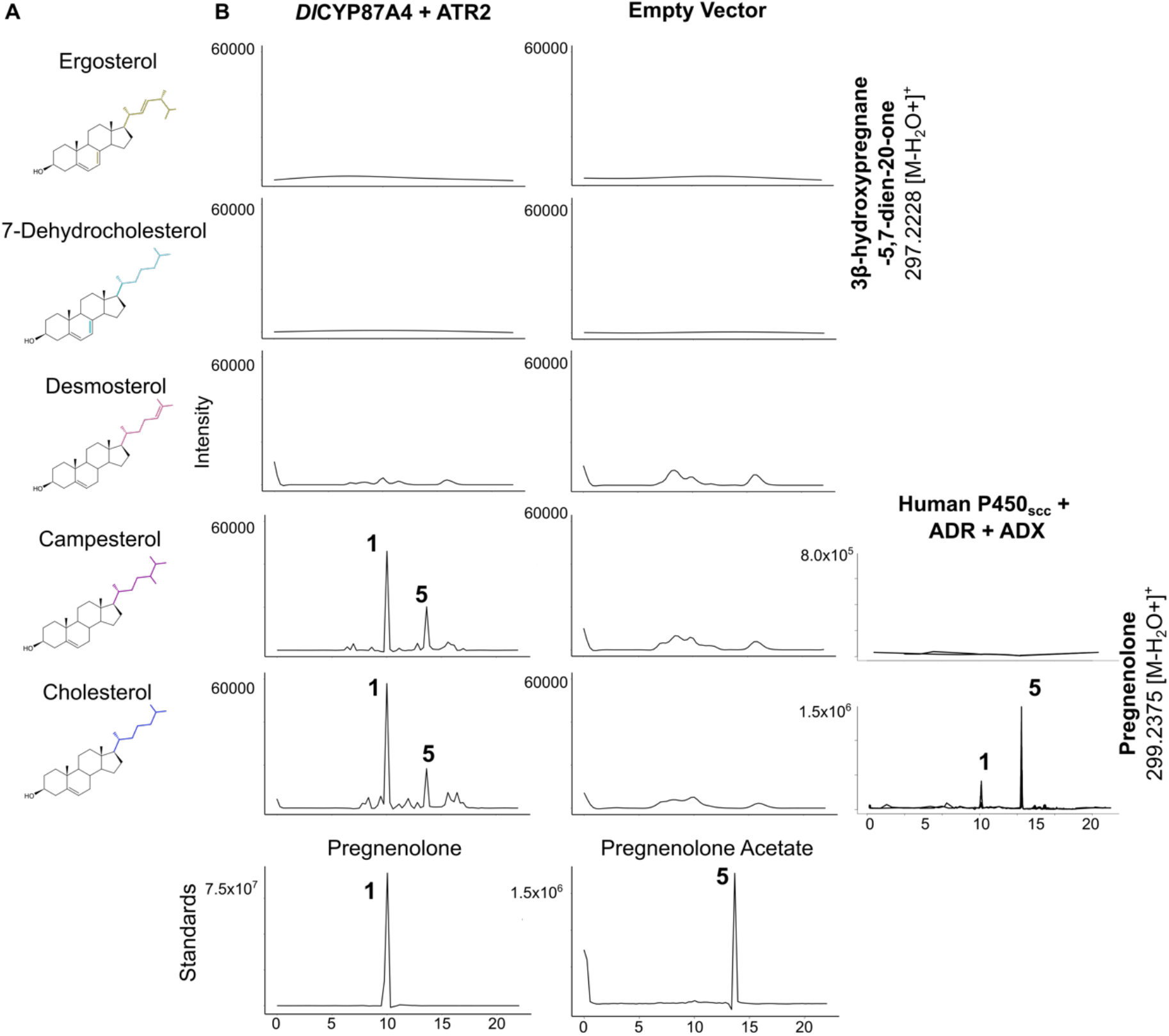
Characterizing the *D. lanata* CYP87A4 in engineered *S. cerevisae* strains containing various sterols. A. Main sterols in each of the yeast strain used in B. B. Extracted ion chromatograms from *in vivo* expression of *Dl*CYP87A4 and ATR in the engineered yeast strains. The human P450_scc_ with its redox partners, adrenodoxin (ADX) and adrenodoxin reductase (ADR) were expressed in campesterol- and cholesterol-producing yeast as a positive control. Authentic standards of pregnenolone and pregnenolone acetate are shown in the bottom panel.

### Neofunctionalization of CYP87A4 unique to *Digitalis*

In order to understand the evolutionary history of the *D. lanata* CYP87A4 enzyme, we constructed a phylogenetic tree with transcript homologous to the *Dl*CYP87A4 from the 1000 transcriptome project (Figure 4)^44^. Four transcripts in the *D. lanata* transcriptome fall into the CYP87A subfamily. *Dl*CYP87A1 and *Dl*CYP87A2 are 72.2 and 74.2% identical to the characterized *Dl*CYP87A4 (Supplementary Table S2). *Dl*CYP87A3 is 97.4% identical at the protein level to the characterized *Dl*CYP87A4 but failed to amplify from the cDNA. It is likely that *Dl*CYP87A1 and *Dl*CYP87A2 represent the canonical enzymes of the CYP87A subfamily whose function is still unknown. Indeed, *Dl*CYP87A1 and *Dl*CYP87A2 are expressed almost constitutively in leaves and roots (Figure 2A). On the other hand, *Dl*CYP87A3 and *Dl*CYP87A4 may originate from duplication and neofunctionalization rendering its sterol cleavage activity. It is important to note that the *Dl*CYP87A4 does not share significant sequence homology to the mammalian P450_scc_. None of the other *Lamiales* in the 1000 transcriptome project had duplicates of the CYP87A subfamily. Species in the *Oenothera* genus also have multiple copies of CYP87A but their function is unclear since these species are not known for producing cardiac glycosides.

**Figure 4:**
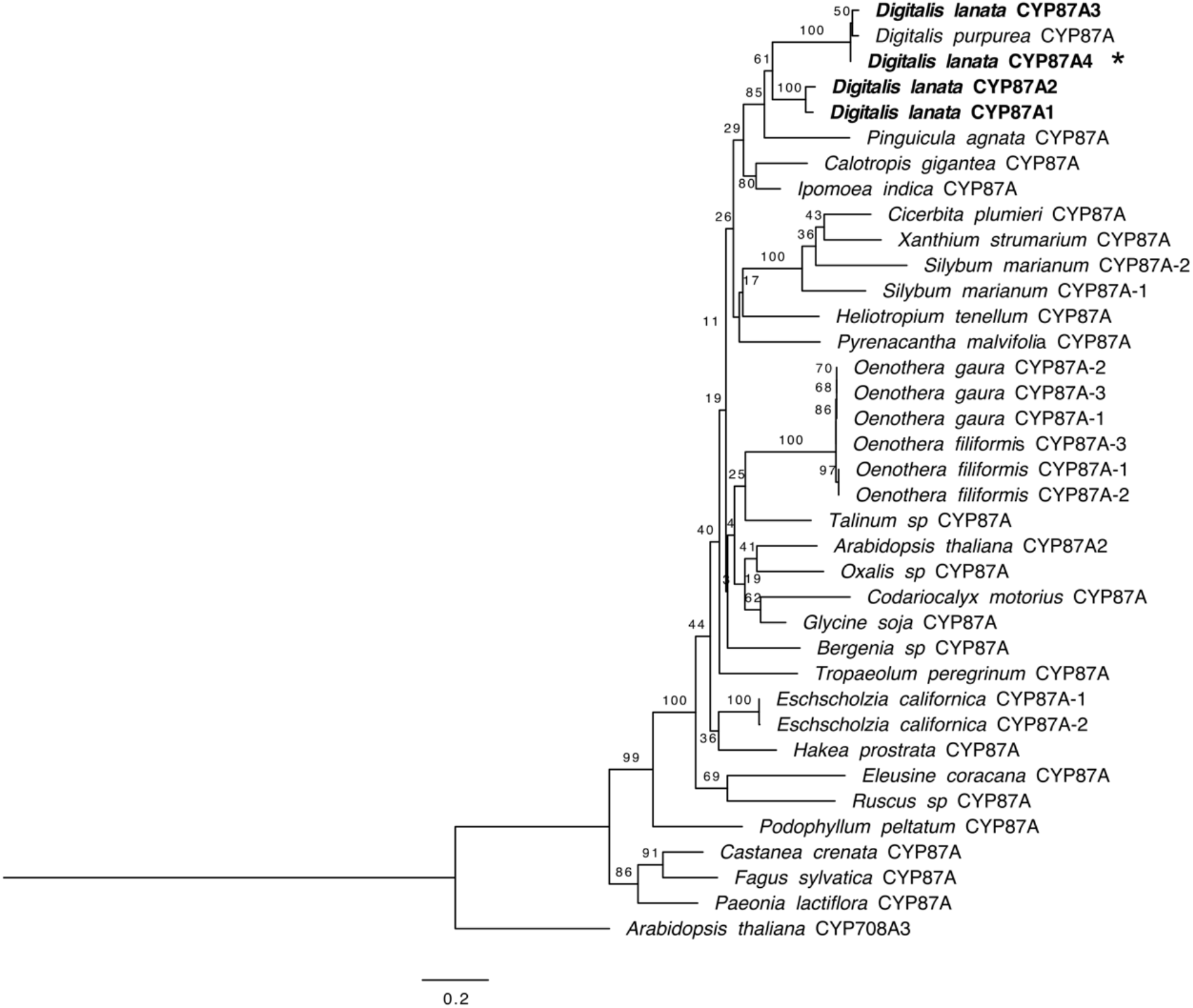
Maximum-likelihood phylogenetic tree of the CYP87A family members from eudicot plants in the one thousand plant transcriptome project. Protein sequences are retrieved from 1,000 plant transcriptome project and filtered for sequences with a start codon, between 400-600 amino acids long, and are 50% identical on the protein level to *Dl*CYP87A4. Bootstrap values from 1000 replicates are shown at each node. Scale bar represents mean number of substitutions per amino acid.

We also included similar transcripts from other plants that produce cardiac glycosides since they likely have an enzyme with a similar sterol cleaving function^10,18^. We searched the publicly available transcriptomes of *Digitalis purpurea, Calotropis gigantea*, and *Asclepias syriaca* for transcripts that were a close match to the *Dl*CYP87A4. *A. syriaca* did not have any transcripts that match over 55% at the protein level with *Dl*CYP87A4. *C. gigantea* had one transcript which matched 69% to *Dl*CYP87A4 (Figure 4, Supplementary Table S2). The *C. gigantea* CYP87A is likely the canonical CYP87A enzyme since there is only one copy and fits well within the canonical CYP87A clade. The *D. purpurea* transcriptome had one transcript that matched 97% to the *Dl*CYP87A4 and was clustered with *Dl*CYP87A3 and *Dl*CYP87A4 on the phylogenetic tree (Figure 4, Supplementary Table S2). Thus, this transcript likely has the same sterol cleaving ability. No canonical CYP87A copy was identified from the *D. purpurea* transcriptome but it is uncertain if it was due to the loss of the gene over time or poor transcriptome quality. Our analysis indicates that the expansion and neofunctionalization of *Dl*CYP87A as P450_scc_ is likely unique to the *Digitalis* species and the sterol cleaving enzymes of *A. syriaca* and *C. gigantea* are still unknown.

## Discussion

We identified and characterized the first and rate-limiting enzyme in the plant cardiac glycoside biosynthetic pathway, P450scc, an enzyme that has long been speculated but never found before. A “cholesterol side-chain-cleaving enzyme” (SCC) that converts cholesterol to pregnenolone in plants was first hypothesized by Pilgrim in 1972, or 50 years ago^24^. Using differential transcriptomic analysis, we showed that a CYP87A family protein acts as the P450scc in Digitalis and has no obvious sequence homology with its mammalian counterpart, CYP11A1. The mammalian P450scc converts cholesterol to pregnenolone, the first and rate-limiting step in steroid hormone biosynthesis ^45^. The *Digitalis* P450_scc_ catalyzes the same reaction in plants but is a completely different enzyme. Additionally, the mammalian P450_scc_ uses adrenodoxin and adrenodoxin reductase as redox partners in order to pass NADPH to the heme center^23^. However, plants do not seem to have these redox proteins and the plant P450_scc_ likely uses a plant cytochrome P450 reductase as its redox partner (Figure 3)^42^. The similarities and differences between the mammalian and plant P450_scc_ indicate that the “cholesterol side-chain-cleaving” activity evolved independently, providing an elegant example of convergent evolution in animals and plants.

The *Dl*P450_scc_ identified is a crucial “gatekeeping” enzyme that connects plant primary metabolism to the secondary metabolism. It channels sterols essential for maintaining cell membrane homeostasis to produce cardiac glycosides, secondary metabolites that play an important role in defense^28^. The *Dl*P450_scc_ is rate-limiting in the cardiac glycoside biosynthetic pathway. This was shown when feeding cholesterol to *Digitalis* embryo only produced a trace amount of pregnenolone whereas administering progesterone increased various pregnane intermediates in the pathway^22^. Furthermore, the fact that pregnenolone is not detected in *Digitalis* but other intermediates in the pathway are detected also support that *Dl*P450_scc_ is a bottleneck enzyme in digoxin biosynthesis (Supplementary Figure S10). Unlike P450_scc_ in animals, *Dl*P450_scc_ is promiscuous as it can catalyze the side-chain-cleaving reaction for both cholesterol and campesterol (Figure 3 Supplementary Figure S12). This is somewhat expected since campesterol is one of the major sterols in plants. Enzyme promiscuity may shape the formation of complex metabolic networks essential for the robustness and resistance of living organisms, especially in plant secondary metabolism^46^. The discovery of *Dl*P450_scc_ as a promiscuous protein for both cholesterol and campesterol will likely end the half-a-century controversy over the sterol precursor for digoxin, although direct *in planta* evidence in *Digitalis* is still needed.

Future work is necessary to understand if the *Dl*P450_scc_ acts by the same catalytic mechanism as the mammalian P450_scc_ which catalyzes three-step sequential oxidations through 22-hydroxylation, 20-hydroxylation, and cleavage between C20 and C22^23^. Docking sterol substrates to the reaction center of a *Dl*P450_scc_ model (Supplementary Figure S13) and previous *in vitro* assay using 20-or 22-hydroxycholesterol for *Dl*P450_scc_ supports this mechanism^22^.

The identification of *Dl*P450_scc_ will enable the study of cardiac glycoside biosynthesis in other plant species such as milkweed (*Asclepias, Calotropis*), wall flower (*Erysimum*), oleander (*Nerium oleander*), lily of the valley (*Convallaria majalis*), just to name a few^47^. Phylogenetic analysis showed that *Calotropis gigantea* may not have a duplicated CYP87A gene (Figure 4) indicating another cytochrome P450 acts as the P450_scc_ in this species, assuming the publicly available *Calotropis* transcriptome is complete. Interestingly, the CYP87A is in the same phylogenetic clade as CYP90B1 that catalyzes the 22*(S)-*hydroxylation of campesterol, one of the three catalytic steps in the sterol side chain cleaving reaction (Supplementary Figure S7)^23^. Thus, cytochrome P450s that fall into this clan, including CYP708A, CYP88A, CYP702A, CYP85A, CYP90, CYP720A, and CYP724A, may all have the potential to acquire the sterol side-chain-cleaving activity. The function of the canonical CYP87A, however, is still unknown. It is likely to oxidize a sterol or a triterpenoid since CYP87D16 from *Maesa lanceolata* oxidizes the C16 of β-amyrin ^48^.

In conclusion, this work identified the rate-limiting, and long-speculated P450_scc_ in *Digitalis* for the biosynthesis of digoxin. It is an important step forward towards complete elucidation of digoxin biosynthetic pathway. The high-quality *D. lanata* transcriptome provides a valuable resource to uncover other digoxin pathway enzymes. This work will also open the door for biomanufacturing novel digoxin analogs with expanded medicinal value either in microbial or plant systems.

## Methods

### Plant material, RNA isolation, and sequencing

*Digitalis lanata* Ehrh. seeds were procured from Strictly Medicinal (Williams, Oregon, USA). These seeds were germinated on the soil mix (57 g triple superphosphate, 85 g calcium hydroxide, 57 g bone meal, 369 g Osmocote (14-14-14), 99 g calcium carbonate, 25 L perlite, 50 L loosened peat and 25 L coarse vermiculite) and maintained in a growth chamber (Invitrogen, Clayton, Missouri, USA) under a light period of 16-hour at 25 °C and relative humidity of 60-80%. The true leaves and whole roots from 5-week-old seedlings were used for RNA isolation. Freshly collected samples were snap-frozen in liquid nitrogen and stored at −80°C before RNA extraction.

Leaf and root tissues from three different seedlings were used to prepare the Illumina sequence library. Total RNA was isolated using the RNeasy Plant Mini Kit (Qiagen,Germantown, MD, USA) following the manufacturer’s protocol. The sequencing library was prepared from total RNA using the TrueSeq Ribo-Zero Plant RNA library prep kit (Illumina, San Diego, CA, USA) that removes ribosomal RNA. A quality check of the library was carried out on an Agilent 2100 bioanalyzer. The library was sequenced using Illumina HiSeq 2500 to generate 100 bp paired-end raw reads.

### Transcriptome assembly, annotation, and differential expression analysis

See supplementary materials and methods.

### Real-time polymerase chain reaction (RT-PCR)

Leaves from 3-4 weeks old *D. lanata* seedlings were used for RNA isolation. Freshly collected samples were snap-frozen in liquid nitrogen and ground into powder. Trizol (Ambion, Austin, TX, USA) reagent was used to isolate total RNA according to manufacture’s protocol. DNA in the sample was removed using the TURBO DNA-*free*™ kit (Invitrogen, Waltham, MA, USA). cDNA was synthesized using the iScript™ cDNA Synthesis Kit (BIO-RAD, Herculese, CA, USA) as per the manufacturer’s instructions. qRT-PCR was carried out using the iTaq Universal SYBR Green Super mix (BIO-RAD, Herculese, CA, USA) with *D. lanata* cDNA, gene specific primers. The following thermocycling conditions were used; denaturation at 95 °C for 3 min, 40 cycles of denaturation at 95 °C for 10 sec, and annealing at 55 °C for 30 sec. A melting curve was generated by increasing the temperature from 55 °C to 95 °C in 0.5 °C increments for 81 cycles. Three biological replicates and two technical replicates were included for each sample. Polyubiquitin 10 (UBQ10) was used as an internal standard. Primers are listed in Supplementary Table S3.

### Gene isolation and Cloning

#### pEAQ system for Nicotiana benthamiana expression

Genes of interest were PCR-amplified from the cDNA of *D. lanata* leaves using either Platinum™ Taq (Invitrogen, Waltham, MA, USA) or Phusion^®^ DNA polymerase (New England Biolabs (NEB), Ipswitch, MA, USA) and cloned into the pEAQ binary vector^49^. The pEAQ plasmid was first digested using the AgeI and XhoI restriction enzymes (NEB, Ipswitch, MA, USA) and the gene of interest was cloned into the linearized pEAQ using the Gibson cloning method ^50^, Genes inserted were sequence verified by Sanger sequencing (Genewiz, South Plainfield, New Jersey, USA).

#### MoClo system for yeast expression

We used the Golden Gate cloning based yeast MoClo system to clone genes of interest for yeast expression ^51,52^. *D. lanata P450*_*scc*_ was PCR-amplified from the pEAQ_*DlP450*_*sc*c_. Human *P450*_*scc*_ along with its redox partners, *ADX* and *ADR*, were amplified from three pYTK001 part plasmids carrying these tree genes. The *Arabidopsis thaliana* redox partner *ATR2* was synthesized by TWIST biosciences (San Francisco, CA, USA). All genes were cloned into the MoClo pYTK001 entry vector using the Esp3I (a high-fidelity analog of BsmBI) restriction enzyme (NEB, Ipswitch, MA, USA). The following point mutations were made in *Dl*CYP87A4 using primers: S123A, A355V, and L357A (See Supplementary Table S3 for primers). Genes cloned were by Sanger sequencing (Genewiz, South Plainfield, New Jersey, USA). Transcription-unit (TU) plasmids were assembled with promoters and terminators from the MoClo kit using high-fidelity BsaI restriction enzyme (NEB, Ipswitch, MA, USA)^51^. TU plasmids constructed were *pTDH3-DlP450*_*scc*_*-tENO1, pCCW12-ATR2-tSSA1, pTDH3-HsP450*_*scc*_*-tENO1, pCCW12-HsADX-tSSA1*, and *pPGK1-HsADR-tADH1*. The transcription units were then used to assemble a multigene plasmid with a 2μ yeast origin of replication. Primers used are listed in Supplementary Table S3. The yeast strains and plasmid constructs used in this study are listed in Supplementary Tables S4 and S5 respectively.

### Tobacco transient expression

#### Agrobacterium transformation

pEAQ plasmids carrying genes of interest were transformed into the *Agrobacterium tumefaciens* strain AGL1 individually by the freeze-thaw method ^53^. The resulting strains were prepared for infiltration using a modified protocol as in Saxena, P. et.al ^54^. Briefly, A single *Agrobacterium* colony containing one of the cloned pEAQ plasmids was inoculated into 5 ml yeast extract broth (YEB) medium (5 g/L tryptone, 1 g/L yeast extract, 2.5 g/L luria broth (Fisher scientific, Waltham, MA, USA), 5 g/L sucrose,0.49 g/L MgSO_4_·7H_2_O) with 50 mg/L kanamycin for pEAQ plasmid selection and 25 mg/L rifampicin for *A. tumefaciens* strain AGL1 selection. The bacterial cultures were grown for 24 hours at 28 °C with shaking at 220 rpm. Afterwards, 0.5 ml of the seed culture was used to inoculate 25 ml of YEB with kanamycin (50mg/L) and rifampicin (25 mg/L) in a shaking flask, and the flasks were grown overnight at 28 °C at 220 rpm shaking. The cultures were pelleted by centrifugation at 3,000g for 15 min, washed once with 10 mL sterile double-distilled water (ddH_2_O), and resuspended in MMA (10 mM MES (2-N-morpholinoethanesulfonic acid), adjusted pH to 5.6 with NaOH, 10 mM MgCl_2_, 100 μM acetosyringone). The individually transformed strains were then pooled together so that the final MMA culture volume was 10 ml and each *A. tumefaciens* strain had a final OD_600_ of 0.4. Then cultures were incubated for 2 to 4 h at 28 °C before infiltrating tobacco leaves.

#### Tobacco infiltration

The pooled *A. tumefaciens* was infiltrated into the underside of four-to six-week-old *Nicotiana benthamiana* new leaves using a needleless plastic syringe and applying gentle pressure. The tobacco plants were grown in 16-hour light, 8-hour dark period at 21 °C with a relative humidity of 60-80 % and photon intensity of 120-150 μmol/m^2^ Three leaves were infiltrated for each experimental set and each set was completed on a single plant. *A. tumefaciens* transformed with pEAQ_*GFP* was infiltrated into a separate plant as the negative control. Plants were maintained in the dark for 12 hours to increase the agrobacterial infection and then shifted to normal light period. The plants were maintained in normal light and growth conditions for an additional four to six days. Once the fluorescence was intense when GFP control leaves were exposed to UV light, all infiltrated leaves were then detached from the petiole, snap-frozen in liquid nitrogen, and ground into a fine powder. Metabolites were extracted by adding 1ml 100% methanol (Fisher scientific, Waltham, MA, USA) and heating at 65°C for 10 minutes. They were then centrifuged at 17,000 g for 10 minutes, filtered through a 0.45 μm filter (VWR, Randor, PA, USA), and stored at −20 °C before LC/MS analysis.

### Yeast *in vivo* expression assay

Cholesterol-producing yeast (strain RH6829), campesterol-producing yeast(strain RH6827), desmosterol-producing yeast (strain RH6828), and 7-dehydrocholesterol-producing yeast (strain RH6826) were kindly provided from the Riezman lab (Supplementary Table S4) ^37^. The wildtype yeast strain BY4741 produces ergosterol natively. Competent cells of these strains were prepared using the Frozen EZ Yeast Transformation II Kit™ (Zymo Research, Irvine, CA, USA) and transformed with a multigene plasmid carrying *DlCYP87A4* and *ATR2* or human *P450scc, ADX*, and *ADR*. Starter cultures were grown in the yeast auxotrophic synthetic dropout (SD-Leu) medium at 30°C overnight and then used to inoculate 25 ml SD-leu medium in a shaking flask with an initial OD_600_ of 0.2-0.4. Samples were harvested at 18 h and pelleted 3,000 g for 5 minutes. Yeast cells were resuspended in 200 μL of TES buffer [50 mM Tris-HCl pH_=_7, 600 mM sorbitol, 10 g/L bovine serum albumin (BSA),1.5 mM β-mercaptoethanol] and homogenized with an equal volume of 0.5 mm glass beads in a BBX24 Bullet Blender^®^ homogenizer (Next Advance, Troy, NY, USA) at setting 8 at 4°C for 4 minutes. 300 μL of TES buffer was added to the lysed cells, and 400-500 μL of the yeast lysate was transferred into a capped glass tube followed by adding 1 mL of chloroform immediately. The sample was vortexed for 1 minute, and the organic phase was transferred into a new glass test tube and dried under a stream of air. The sample was resuspended in 100 μL 100% methanol, centrifuged at 17,000 g for 10 min, and the supernatant was transferred into a glass LC/MS tube and stored at −20 °C until use.

### LC/MS analysis for pregnane intermediates in the digoxin pathway

*Digitalis lanata* and tobacco extract samples were analyzed as previously described and yeast lysate samples were run as previously described with the following modifications ^55^. Samples were analyzed using LC coupled with a tandem high-resolution MS instrument, a Thermo Scientific Q-Exactive Focus™ (Fisher scientific, Waltham, MA, USA). A Waters XSelect CSH™ C18 HPLC column (Waters, Milford, MA, USA) with a particle size of 3.5 μm, an internal diameter of 2.1 mm and a length of 150 mm was used for the separation. For analyzing primary cardenolides including lanatoside A, B, C, and E, and cardenolide pathway intermediates, including pregnenolone, progesterone, 5-*β*-pregnane-3,20-dione, 3*β*-hydroxy-pregnane-20-one, 3*β*,14*β*-dihydroxy-5*β*-pregnane-20-one, and 3*β*,14*β*,21-trihydroxy-5*β*-pregnane-20-one from plant samples the LC/MS protocol is the same as previously described ^56^. For analyzing pregnenolone and pregnenolone acetate from yeast samples, the following protocol was developed. The mobile phases were water with 0.1% formic acid (mobile phase A) and acetonitrile with 0.1% formic acid (mobile phase B) operated at a flow rate of 200 μL min^-1^. Gradient elution started with 40% mobile phase B for 2 min followed by a linear gradient 40∼95% B from 2-12 min, held for 5 min and brought back to initial conditions of 40% mobile phase B for 1 min. Sample injection volume was 20 *μ*L. A full-scan data-dependent MS^2^ was used with an inclusion list of the precursor ion mass which included a scan range *m/z* 100-1200, and all other settings as previously described^56^. The mass spectrometer was regularly calibrated in both positive and negative mode with a solution of compounds of known mass. Qualitative analysis was preformed using the XCalibur™ software and chromatograms and spectra generated in R using the XCMS and Spectra packages ^57,58^.

### Gas chromatography coupled mass spectrometry (GC/MS) analysis for cholesterol and phytosterol quantification

For steroid analysis, *D. lanata* leaf and root samples were prepared as described previously by Itkin, M. et. al ^59^. Yeast samples were prepared by lysing OD_600=_1 of cells. Yeast cells were pelleted at 500g for 4 minutes, resuspended in 200 μL of TES buffer, and homogenized with equal volume of 0.5 mm glass beads in a BBX24 Bullet Blender^®^ homogenizer (Next Advance, Troy, NY, USA) at setting 8 at 4°C for 4 minutes. 300 μL of TES buffer was added to the lysed cells, and 400-500 μL of the yeast lysate was transferred into a capped glass tube and resuspended in 6 mL chloroform/methanol mix (2:1, v/v). Samples were heated at 75°C for 60 min. Solvent was dried under a stream of air and resuspended in 2 mL of 6% (w/v) KOH in 100% methanol. Samples were saponified by heating at 90 °C for 60 minutes. They were then cooled to room temperature and 1.5 mL hexane and 1.5 mL ddH_2_O was added followed by vigorously shaken by hand for 20 sec. The organic hexane phase was separated by centrifugation at 3,000g for 2 min and 1 mL was transferred to a new glass test tube. The sample was dried under a stream of air and resuspended in 50μL of N-methyl-N-(trimethylsilyl) trifuoroacetamide (MSTFA) (Fisher scientific, Waltham, MA, USA), votexed to mix for 20 sec, and transferred to a glass insert inside a GC glass vial for derivatization.

Samples were analyzed on the Thermo Scientific™ Q-Exactive™ GC Orbitrap™ (Fisher scientific, Waltham, MA, USA). Samples were injected into a Thermo Scientific TraceGOLD TG-5SILMS column that was 30 m long, had a 0.2 mm inner diameter and 0.25 film thickness. Inlet temperature was set at 280°C. Helium was used as carrier gas at a flow rate of 1.2 mL/min. The thermal gradient started at 170°C and held for 1.5 min, then ramped to 280°C at 37°C/min, further ramped to 300°C at 1.5°C/min, and finally held at 300°C for 5.0 min. Eluents were ionized by electron impact ionization at 70 eV. A filament delay of 6 min was used. High resolution EI fragment spectra were acquired using 60,000 resolution (FWHM at m/z 200).

### Phylogenetic analysis

Cytochrome P450s for the cytochrome P450 tree were retrieved by searching the transcriptomes for transcripts that had proteins containing the Pfam domain PF00067 (CYPs) and were between 400-600 amino acids in length. *Arabidopsis thaliana* sequences for the cytochrome P450 tree (Supplementary Figure S7) were retrieved from the *Arabidopsis* CYP database (Paquette S.M. *et al*, 2000).

Transcriptome sequences used for the CYP87A tree were retrieved by BLASTing the *Dl*CPY87A4 transcript against the 1000 Plant Transcriptome (1KP) database using tBLASTx (Leebens-Mack J.H. *et*.*al*. 2019, Carpenter E.J. *et*.*al. 2019*). Sequences were filtered and only those that were within 400-600 amino acids long and had a start codon were retained. *A. thaliana* cDNA from CYP87A2 was used as a reference and CYP708 was used as an outgroup.

All trees were constructed by aligning protein sequences using MAFFT ^60^. Aligned sequences were trimmed using trimAl ^61^. Phylogenetic tree was constructed using RAxML-NG (1.0.1) with the all-in-one Maximum likelihood (ML) tree search and slow bootstrapping with 1,000 replicates ^62^.

### Protein Modeling and Docking

A protein model for the *D. lanata* P450_scc_ (*Dl*CYP87A4) was generated using Alphafold2 through the ColabFold platform v1.4 ^63,64^. The MSA mode used was MMseqs2 and all other parameters were the default parameters ^65^. Five models were generated and model 3 was used for further analysis based on Predicted Aligned Error (PAE) and predicted local distance difference test (pLDDT) scores. Docking of campesterol and cholesterol was performed using Chimera version 1.16 and Autodock Vina version 1.1.2 ^66,67^.

## Acknowledgement

The authors are grateful to Dr. Howard Riezman at University of Geneva for providing the engineered sterol-producing yeast strains. We also thank Rian Hammond for providing plasmids with genes encoding the human P450_scc_, adrenodoxin, and adrenodoxin reductase. We are also thankful to Dr. Valerie Freichs for assistance with chromatography work and Dr. Donald Yergeau for RNA-seq at University at Buffalo.

## Author contributions

E.C., B.R.G., and Z.Q.W. designed research; E.C., B.R.G., and I.R. carried out experiments; E.C., B.R.G, I.R., and Z.Q.W. analyzed data; E.C., B.R.G, I.R. and Z.Q.W. wrote the paper.

## Data Availability

Data is available upon request from the authors.

