## Supplementary Information for "The P450 Sterol Side Chain Cleaving Enzyme (P450_scc_) for Digoxin Biosynthesis in the Foxglove Plant Belongs to the CYP87A Family"

| **Supplementary Materials and Methods** | **Page** |
|---|---|
| *De novo assembly of transcriptome* | 3 |
| *Transcriptome annotation and functional classification* | 3 |
| *Identification of differentially expressed transcripts* | 4 |
| *Identification of transcription factor and protein kinase families* | 4 |
| *Identification of simple sequence repeats (SSRs)* | 4 |
| *BUSCO analysis* | 5 |
| **Supplementary Results and Discussion** | |
| *Transcriptome sequencing and its de novo assembly* | 6 |
| *Transcriptome annotation* | 6 |
| *Gene ontology (GO) classification* | 8 |
| *Functional characterization using KEGG* | 8 |
| *Protein kinase, transcription regulator, and transcription factor families* | 12 |
| *Simple sequence repeats (SSRs)* | 14 |
| **Supplementary Figures and Tables** | |
| *Figure S1. Statistical analysis of the assembled transcriptome* | 7 |
| *Figure S2. Functional classification of the unigenes from the transcriptome* | 9 |
| *Figure S3. KEGG pathway annotation* | 11 |

*Figure S4. PKs, TFs, TRs in the D. lanata transcriptome* 13

*Table S1. Top 20 simple sequence repeats (SSRs) in a total of 22,549 D. lanata SSRs* 15

*Figure S5. GC/MS analysis of steroids in the leaves of D. lanata* 16

*Figure S6. Differential expression of terpenoid biosynthetic pathways in leaves vs. roots.* 17

*Figure S7. Phylogenetic tree with D. lanata and Arabidopsis thaliana cytochrome P450s* 18

*Figure S8. Differentially expressed cytochrome P450s in the leaves and roots of D. lanata* 19

*Figure S9. MS spectra of pathway intermediates in tobacco transiently expressing digoxin biosynthetic pathway genes* 20

*Figure S10. Identifying putative digoxin pathway intermediates in D. lanata* 21

*Figure S11. GC/MS analysis of sterol contents in engineered yeast strains* 22

*Figure S12. Preliminary experiment with campesterol-producing yeast expressing DlCYP87A4* 23

*Figure S13. Docking of campesterol and cholesterol into the active site of DlCYP87A4 model* 24

*Table S2. DlCYP87A4 with closely related cytochrome P450s in various plant species* 25

*Table S3. Primers used in this study* 26

*Table S4. Yeast strains used in this study* 27

*Table S5. Constructs used in this study* 28

**Reference** 29

### Supplementary Materials and Methods

#### *De novo assembly of transcriptome*

FastQC (v0.11.5) was used to check the next-generation sequencing data [1]. Then, the Trimmomatic tool embedded in the Trinity transcriptome assembler was used to remove Illumina sequencing adaptors, low-quality leading and trailing bases (below quality 5), cutting a sliding window of size 4 bp when their average quality drops below 5 and also filtering out reads below 25 bp. *De novo* assembly of a single transcriptome (leaf and root) from the raw reads was done using the Trinity assembler program (v2.8.4) at the default k-mer value of 25 [2]. Bowtie2 (v2.3.4.3) was used to map raw reads to the assembled transcripts to ascertain completeness [3]. The overall quality of the transcriptome was ascertained using the N50 and ExN50 values calculated using the Trinity pipeline.

#### *Transcriptome annotation and functional classification*

Firstly, the open reading frames (ORFs) and their corresponding protein sequences were identified from the transcripts of the transcriptome using the TransDecoder tool in the Trinity pipeline. Identification of the unigenes (unique ORFs) was carried out using CD-HIT-EST (v4.6.8) with an identity threshold of 99%, a word length of 10 (the size of the sequence used in each cycle of comparison), and with the mode to cluster the most similar sequences [4]. The transcriptome was annotated using the BLASTx and BLASTp tools [5]. NCBI non-redundant protein database [6], UniProt [7], and Pfam [8] were used to search for protein homologs in the ORFs with a cut-off e-value of 1e-5. InterProScan (v5.32-71.0) was used to scan the ORFs for various protein signatures [9]. Databases, namely CATH-Gene3D, CDD, HAMAP, MobiDB, PANTHER, Pfam, PIRSF, PRINTS, ProDom, PROSITE, SMART, SUPERFAMILY, SFLD, and TIGRFAMs were used to identify domains or motifs of proteins represented by the ORFs through InterProScan. GO terms assigned by InterProScan were used by WEGO (v2.0) for the GO annotation of ORFs [10].

KEGG (Kyoto Encyclopedia of Genes and Genomes), GhostKOALA (KEGG orthology and links annotation), and KEGG automated annotation service (KAAS) were used to assign KEGG-orthology (KO annotation) to ORFs. Among the KO annotations, KAAS was given precedence as it identified KO based on a comparison with related species, whereas the GhostKOALA searches the prokaryotic, eukaryotic, and viral sequences. The amino acid unigene sequences were BLASTed against the genes of selected organisms. In our case, we BLASTed against plants including *Brassica napus* (rape), *Citrus sinensis* (Valencia orange), *Theobroma cacao* (cacao), *Gossypium raimondii*, *Gossypium hirsutum* (upland cotton), *Glycine max* (soybean), *Medicago truncatula* (barrel medic), *Rosa chinensis* (China rose), *Vitis vinifera* (wine grape), *Solanum lycopersicum* (tomato), *Nicotiana tabacum* (common tobacco), *Olea europaea* var. sylvestris (wild olive), *Helianthus annuus* (common sunflower), *Oryza sativa* japonica (Japanese rice) (RefSeq), and *Zea mays* (maize) based on the NR BLASTx search results of the total ORFs. A few mammals including *Rattus norvegicus* (rat), *Bos taurus* (cow) and *Homo sapiens* (human), fish *Danio rerio* (zebrafish), insect *Drosophila melanogaster* (fruit fly), and nematode *Caenorhabditis elegans* were considered because they are well studied in the genetic aspect and contain the steroid biosynthesis pathway genes which are of central interest in the current study. A few fungi, such as *Saccharomyces cerevisiae* (budding yeast), *Ashbya gossypii* (Eremothecium gossypii), *Candida albicans*, *Schizosaccharomyces pombe* (fission yeast), and *Encephalitozoon cuniculi* were included to represent majority groups of fungi. Protists such as *Entamoeba histolytica*, *Plasmodium falciparum* 3D7, and *Cryptosporidium hominis*, and prokaryotes such as *Escherichia coli* K-12 MG1655, *Neisseria meningitidis* MC58 (serogroup B), *Helicobacter pylori* 26695,

*Bacillus subtilis* subsp. subtilis 168, *Lactococcus lactis* subsp. lactis Il1403, *Mycoplasma genitalium* G37, *Mycobacterium tuberculosis* H37Rv, *Synechocystis* sp. PCC 6803, *Aquifex aeolicus*, *Methanocaldococcus jannaschii*, and *Aeropyrum pernix* were used for the same reason as mentioned above. A total of 859,009 KEGG sequences were used in the bi-directional best hit method of KAAS BLAST functional annotation of unigenes.

The Reconstruct Pathway tool of KEGG was used to process the KO annotations and further classify the ORFs based on KEGG metabolic pathways, BRITE (hierarchical classifications of biological entities), and modules (functional units of pathways).

#### *Identification of differentially expressed transcripts*

Trinity pipeline was used for differential expression (DE) analysis. As a prerequisite to DE analysis, abundance estimation of the transcripts was done using the alignment-based abundance estimation method, namely "RNA-Seq by Expectation-Maximization" (RSEM) [11]. The abundance data of all the 317,983 transcripts was represented as transcripts per million (TPM) and fragments per kilobase of transcript per million mapped reads (FPKM). A raw counts matrix and a normalized expression matrix were then generated, containing expression data for each replicate of each sample. A "trimmed mean of M values" (TMM) normalization method was used to do cross-sample normalization [12]. Further, the Trinity script using edgeR, a Bioconductor package, was used to conduct the DE analysis to obtain log fold change (logFC), log counts per million (logCPM), p-value, and false discovery rate (FDR) for each transcript. Finally, the expression level of each transcript was studied as TPM where the normalization is first done for transcript length and then for sequence depth to obtain an expression unit that can span different samples and replicates of an experiment [13].

Further, the expression of selected transcripts was studied by plotting the expression data as a hierarchal tree using Multiple Experiment Wiewer (v4.9). Thus, the expression patterns of the various transcripts in relation to each other was observed and co-expressed transcripts were studied as clusters.

#### *Identification of transcription factor and protein kinase families*

The iTAK (v1.7a) stand-alone tool was used to identify plant transcription factors (TFs), transcription regulators (TRs), and protein kinases (PKs) from the protein sequences obtained from the transcriptome via Transdecoder. TFs, TRs, and PKs were further individually classified internally by iTAK to their respective gene families. The documentation and cataloging of the various TFs and TRs were based on the consensus rules drafted from the PlnTFDB database [14] and the PlantTFDB portal [15]. The database entries of iTAk are well-curated to ensure accuracy. Plant PKs are identified based on protein kinase domains (Pfam domains PF00069 and PF07714). The identified PK hits are further classified internally by iTAK using the respective protein kinase Hidden Markov Models (HMMs) [16].

#### *Identification of simple sequence repeats (SSRs)*

The SSR motifs were identified among the transcripts of the leaf and root transcriptome using the MISA stand-alone PERL tool [17]. The parameters used to define the microsatellites were a minimum of 6 repeated for a unit size of 2 nt, a minimum of four repeats for the unit size of 6 nt, a minimum of five repeated for the unit sizes of 3 nt, 4 nt and 5 nt. Mononucleotide motifs were not considered in the current analysis due to the chance of homopolymer tail artifacts formed during sequencing. Simple repeat motifs comprising one SSR and compound repeat motifs containing two or more SSRs with a maximum of 100 nt interrupting them were considered.

#### *BUSCO Analysis*

Completeness of the transcriptome was assessed using BUSCO [18]. Analysis was performed in transcriptome mode using the eudicots_odb10 database.

### Supplementary Results and Discussion

#### *Transcriptome sequencing and its de novo assembly*

The transcriptome sequencing of leaf and root samples provided a total of 173,448,870 raw reads with an average length of 100 bp. After quality assessment using FastQC and read trimming by Trimmomatic, 173,445,956 high-quality reads were used for transcriptome assembly. A gentler trimming strategy (PHRED = 5) was employed to remove only the lowest-quality bases, thus retaining the shorter and lesser expressed transcripts, which were vulnerable to loss in case of a harsh trimming [19]. A total of 310,473,283 bp were assembled into 317,983 transcripts by the Trinity transcriptome assembler containing 183,152 Trinity genes (distinct groups of transcripts identified by Trinity assembler which contain sequences greatly similar to each other and are considered as isoforms). The average size of transcripts was 976 bp for the entire transcriptome. The size distribution of the total assembled transcripts in the transcriptome is consistent with already existing plant transcriptomes such as *Salvia miltiorrhiza* [20], *Withania somnifera* [21], *Chrysanthemum morifolium* [22], *Calotropis procera* [23], *Persea americana* Mill. (Avocado), *Macadamia integrifolia* L. (macadamia), and *Mangifera indica* L. (mango) [24] (Figure S1A). The large number of transcripts whose sizes were greater than 3,500 bp was attributed to the overzealous production of long isoforms by the Trinity assembler (Figure 1A).

Raw reads mapped onto the transcriptome assembly using Bowtie2 (v2.3.4.3) showed 99.36% alignment indicating an excellent quality of the assembled transcriptome [25]. The transcriptome had an overall N50 value of 1,712 bp. Plotting the ExN50 value against varying levels of cumulative transcript expression (Ex) identified a saturation point of the assembly at 88% of the total expression, giving an improved E88N50 of 1,990 bp and reducing the effective transcripts, or transcripts contributing to the saturation point, count to 70,502 (Figure 1B). According to Trinity, the ExN50 peak begins to shift towards ~90% as the read depth increases. Therefore, in this case, since the ExN50 peaks at 88%, the read depth is considered sufficient. This suggests that the assembled transcriptome has a saturation of full-length reconstructed transcripts on account of its read depth (https://github.com/trinityrnaseq/trinityrnaseq/wiki/Transcriptome-Contig-Nx-and-ExN50-stats).

#### *Transcriptome annotation*

Multiple open reading frames (ORFs) were observed in most of the transcripts in the transcriptome, but a good number of them were less than 100 amino acids (300 bp) likely resulting from artifacts of assembly. A total of 190,755 ORFs were obtained from the transcripts using the "TransDecoder.LongOrfs" tool, all the ORFs thus found presented a minimum of 100 amino acids (default for the TransDecoder.LongOrfs tool), among them 121,298 (63.59%) had a methionine start codon and a stop codon, therefore considered complete.

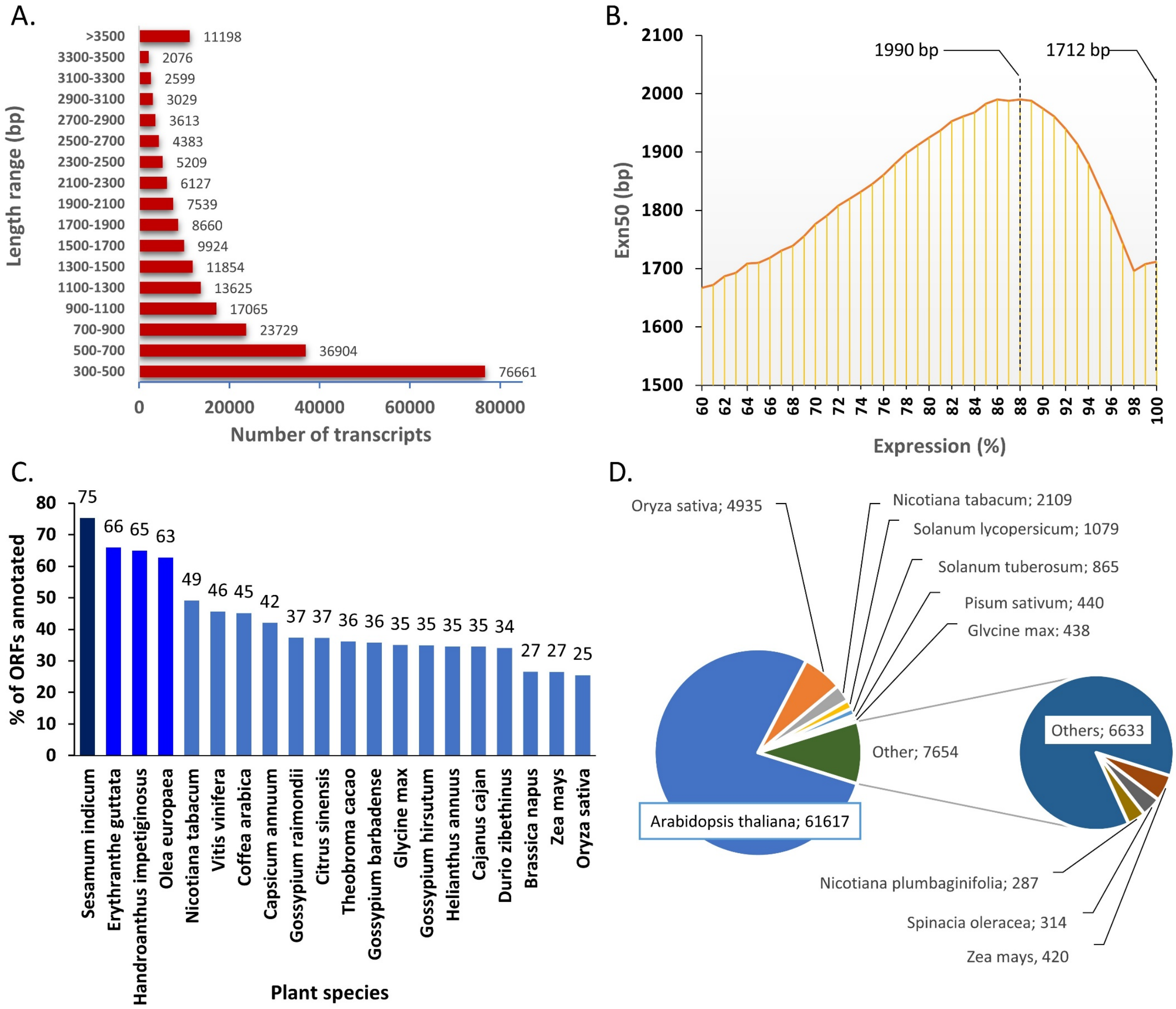


**Figure S1**. Statistical analysis of the assembled transcriptome. A) Transcriptome assembly data showing the size distribution of transcripts (N50 = 1,712 bp). B) Expression-dependent N50 (ExN50) calculated as the N50 of the top X% of the transcript expression levels (Ex). ExN50 of 1,990 bp at the top 88% expression level and traditional N50 of 1,712 bp at 100% expression level are marked. C) Percentage of ORFs annotated by various plant species using BLASTx of the NCBI Non-redundant protein database. D) Number of ORFs annotated by various plant species using BLASTx of the Swiss-Prot non-redundant protein database.

The total number of 5' partials (33,774), or ORFs with a start codon but no stop codon, were a little more than double of the 3' partials (15,036), this is usually seen if poly(A) enrichment is used in the library preparation process [26], but in our case, we used the TrueSeq Ribo-Zero Plant RNA library prep kit with Ribo-Zero ribosomal RNA reduction chemistry and still found similar results. Also, 20,647 sequences were considered internal since they were both 5' and 3' partials. The 190,755 ORFs were annotated with NCBI non-redundant protein database (NR), and 161,326 of the total ORFs were annotated using BLASTx. Taxonomic analysis of the BLAST results revealed that 84.57% of the ORFs matched previously annotated genes, out of which 86.71% represent eukaryotic genes; among the eukaryotic genes, 92.64% represent plant genes.

About 12.88% of the total annotated ORFs represent bacterial genes, and only 0.39% represent archaea. Prominent plant species whose genes are homologous to the annotated ORFs are *Sesamum indicum* (Pedaliaceae), *Olea europaea* (Oleaceae), *Erythranthe guttata* (Phrymaceae), and *Handroanthus impetiginosus* (Bignoniaceae), all belonging to the order of Lamiales, of which *Sesamum indicum* showed a maximum coverage of 75.26% (Figure 1C). UniProt non-redundant curated proteins database (Swiss-Prot) annotated 92,133 (48.30%) ORFs, among which 79,137 (85.89%) were against plant genes. Among the plants, *Arabidopsis thaliana* rendered a maximum annotation of 61,617 (66.88%) ORFs, followed by *Oryza sativa* (5.36%), *Nicotiana tabacum* (2.29%), *Solanum lycopersicum* (1.17%), and *Solanum tuberosum* (0.94%) (Figure 1D).

CD-HIT (v4.6.8) was used to identify 113,221 unigenes with an identity threshold of 99% out of the 190,755 ORFs. Most of the sequences lost are repeats of the same sequences, either complete or partial. The higher threshold of 99% was used so that alleles that have considerable (>1%) sequence variation are retained.

#### *Gene ontology (GO) classification*

Among the total unigenes, 42,724 (37.7%) were linked to gene ontology (GO) terms. 42,724 (44.2%) of these unigenes were classified in the biological process, 11,458 (10.1%) were in the cellular component, and 36,574 (32.3%) were in the molecular function category (Figure S2A). Unigenes in the biological process primarily belong to various metabolic processes, including cellular, organic substance, primary, and nitrogen compound metabolic processes. The majority of unigenes in the cellular component class belonged primarily to the cell part, intracellular part, organelle, and membrane subcategories. Unigenes in the molecular function group were primarily classified in the binding, catalytic activity, heterocyclic compound binding, and ion binding subgroups. A total of 379 GO accessions were observed in the overall GO classification.

#### *Functional characterization using KEGG*

The reconstruct pathway tool of the KEGG mapper analysis annotated 5,683 unigenes [27]. Among them were 3,922 KEGG orthologs (KO) and 1,983 enzymes. Genes and proteins were also identified under the general categories of metabolism, genetic information processing, and signal and cellular processing (Figure 2B). This was completed based on KEGG pathway maps, BRITE hierarchies, and KEGG modules.

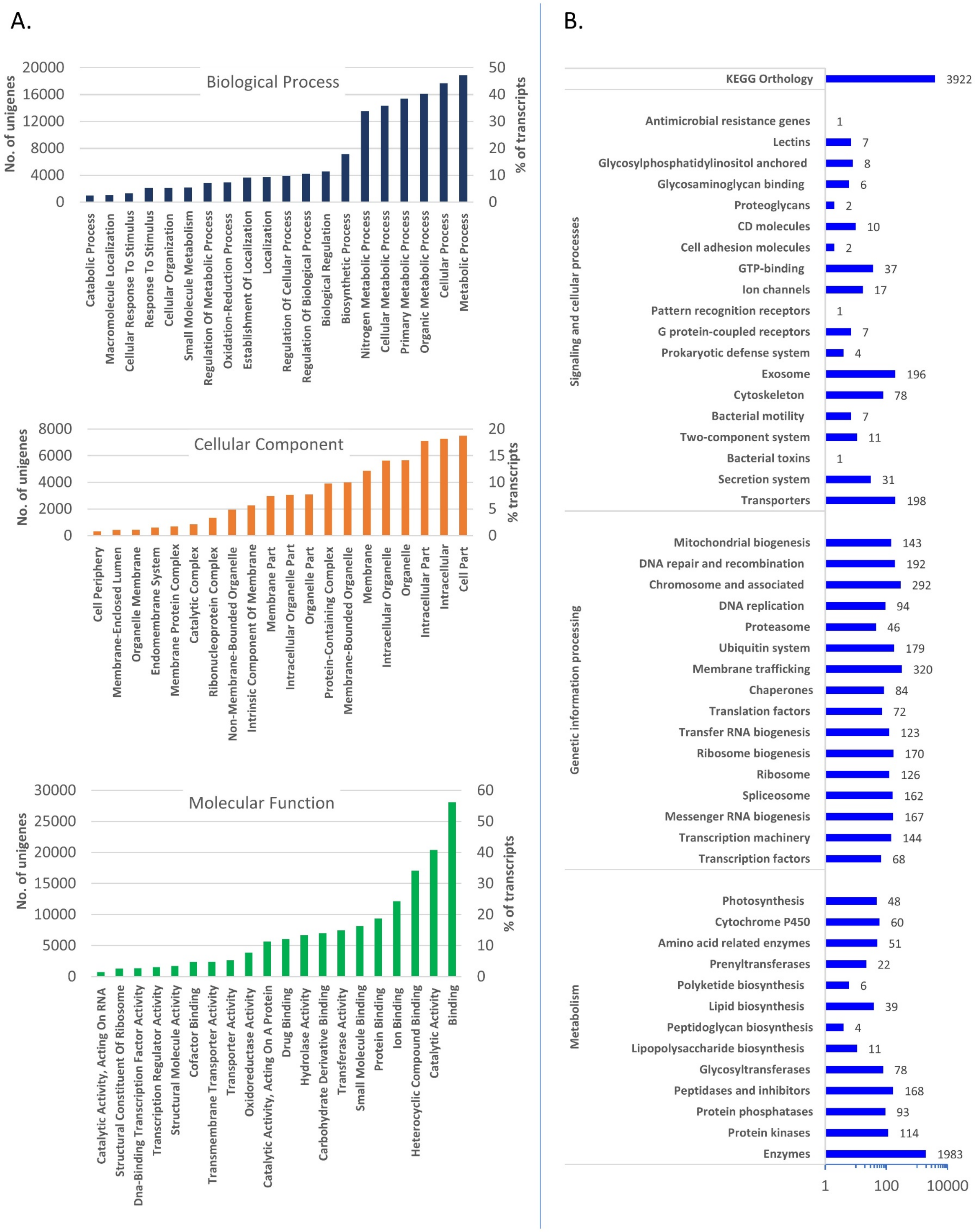


**Figure S2.** Functional classification of the unigenes from the transcriptome. A) Unigenes were plotted as numbers and percentages of unigenes matching various GO categories. B) KEGG mapping against BRITE hierarchies of unigenes classified into various biological processes. The number are various KOs (KEGG orthologs) represented by unigenes.

Although KEGG mapping against BRITE hierarchies is a process similar to GO enrichment, KEGG BRITE currently has 53 classification systems, compared to only three in GO (Figure S3A)[27]. The KEGG mapper revealed representations of unigenes in 412 KEGG pathways, of which 84 modules were complete. Among the complete modules were the terpenoid backbone biosynthesis pathways, including the mevalonate pathway and the non-mevalonate pathway, as well as the mono-, sesqui-, and di-terpenoid biosynthetic pathways (Figure S3B). It also identified partial modules in sterol biosynthesis, such as cholesterol biosynthesis (8 out of 10 enzymes) (Figure S3B). Overall, there were 93 enzymes identified in the metabolism of terpenoids and polyketides.

A.

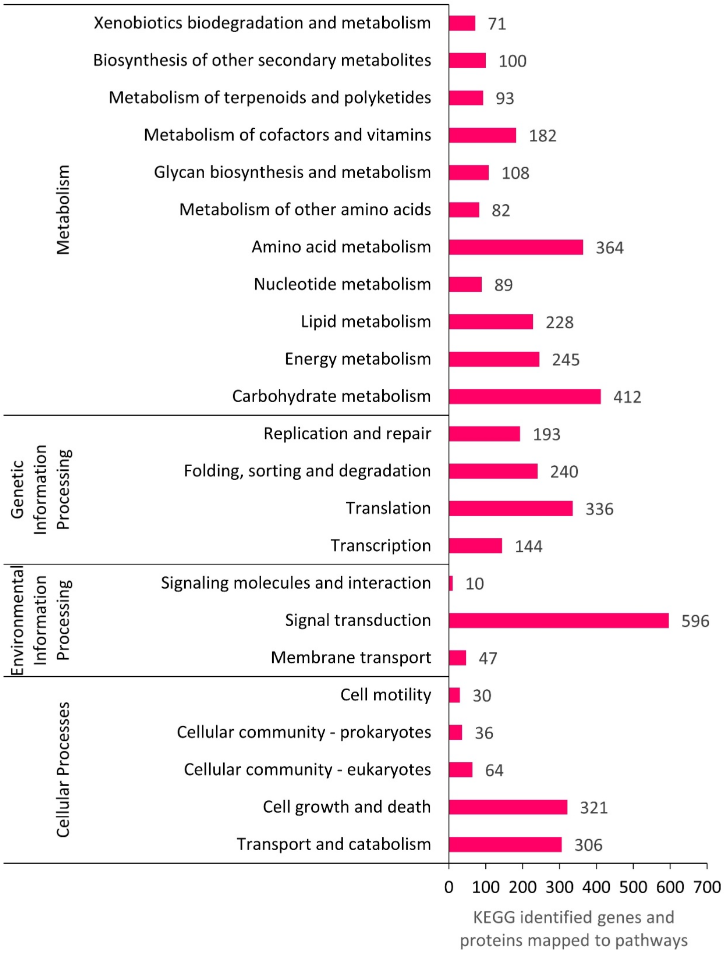


B.

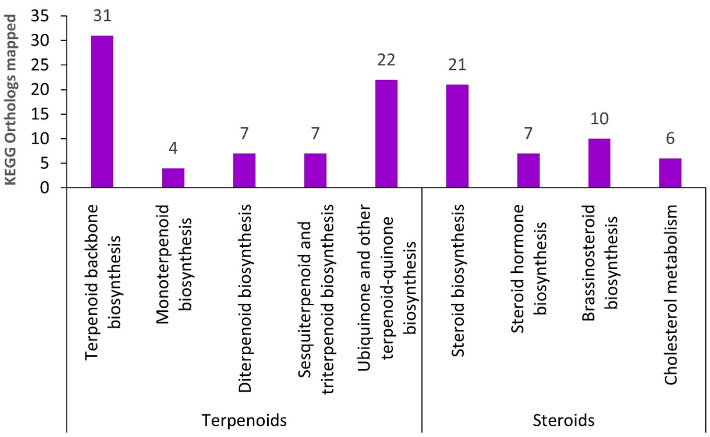


**Figure S3.** KEGG pathway annotation. A) KEGG pathway classification map. Unigenes were allocated biological pathways based on their KO (KEGG ortholog) annotations. The values represent the number of KOs answered by unigenes. B) KEGG analysis showing that KEGG orthologs (KO) involving in terpenoid and steroid pathways are represented by unigenes in this study.

***Protein kinase, transcription regulator, and transcription factor families***

Protein kinases (PKs), transcription regulators (TRs), and transcription factors (TFs) play a key role in plant development and response to environmental stimuli. Much is unknown, especially in the context of secondary metabolism, regarding the role of PKs, TRs, and TFs. Since *D. lanata* has a unique cardenolide pathway, identifying its PKs, TRs, and TFs would lay the foundation for further studies to understand the molecular mechanism of how stresses and environmental stimuli induce the cardenolide pathway.

In the *D. lanata* transcriptome, 126 PK sub-families were identified, of which the RLK-Pelle_DLSV PK sub-family had a maximum representation of 282 unigenes. Also, the RLK-Pelle family seems to be the major family of PK in *D. lanata,* many of the proteins in this family are receptor-like kinases, and the rest are cytoplasmic kinases that lost their extracellular domains [28]. A list of all the unigenes representing various PKs is found in Supplementary file 2 (PK).txt.

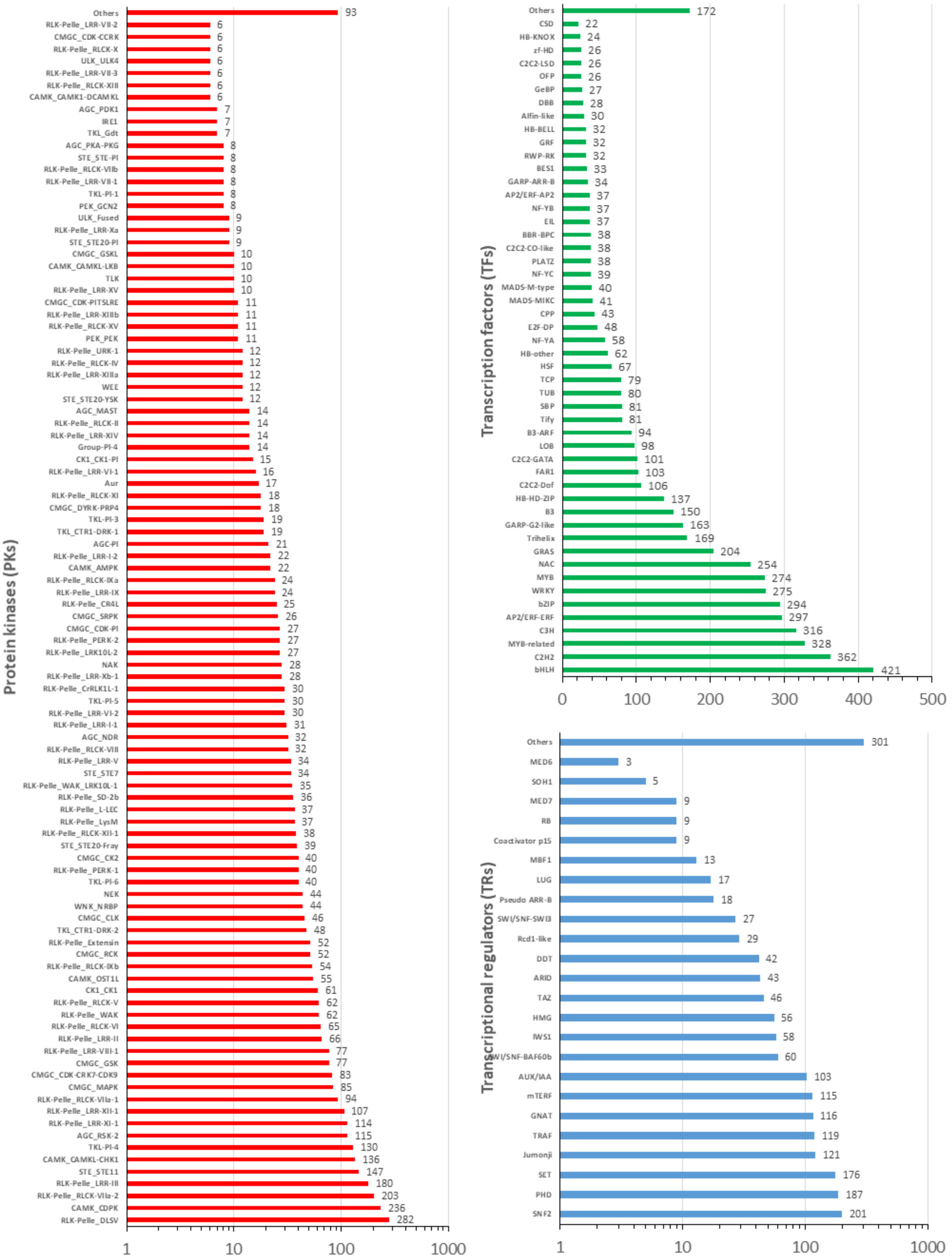


**Figure S4**. Protein kinases (PKs), transcription factors (TFs), and transcription regulators (TRs) in the *D. lanata* transcriptome.

1,883 unigenes were identified as TRs*,* of which 201 unigenes were in the SNF2 (Figure S4A). The SNF2 family plays essential roles in cellular processes such as transcriptional regulation, maintenance of chromosome stability during mitosis, and DNA damage repair [29]. 5,634 unigenes are assigned to 66 TF families, of which the top representation was the bHLH family represented by 421 unigenes (Figure S4B). The basic helix-loop-helix (bHLH) is the most general class of TFs in eukaryotes. bHLH is commonly involved in plant growth and metabolism, especially in photomorphogenesis, light signal transduction, and secondary metabolism. It also plays a vital role in stress response [30]. A list of all the unigenes representing various TRs and TFs is available in “Supplementary file 3 (TR TF).txt”.

#### *Simple sequence repeats (SSRs)*

*De novo* assembled transcriptomes have proven to be a useful tool for molecular marker development [31]. Such markers are predominantly simple sequence repeats (SSRs), also called microsatellites. SSR markers identified from a transcriptome are termed expressed sequence tag-simple sequence repeat (EST-SSR) markers. Although usually low levels of polymorphism are detected in EST-SSRs compared with genomic SSRs, EST-SSRs can be successfully used for various purposes, and they may prove superior to genomic SSR markers for diversity estimation and transferability [32]. These EST-SSRs play a vital role in genetic diversity analysis and plant identification. These SSRs are short repeat sequences with unit sizes ranging from 1 nt to 7 nt. They may also contain more than one motif with overlapping nucleotides. A total of 22,549 SSRs were identified in the *D. lanata* transcriptome. The number of transcripts containing more than one SSR was 2,520, and the number found in compound formations was 1,520. Among the 2-nt, 3-nt, 4-nt, 5-nt, and 6-nt SSR motifs, the total 2-nt SSRs were predominant (12,530), followed by 3-nt SSRs (7,490). The lowest was the 4-nt (673). 5-nt (912), and 6-nt (944) motifs were almost equal. Most 2-nt SSR motifs were in the AG/CT, AT/AT, and AC/GT categories. The 3-nt SSR motifs were predominantly AAG/CTT, AAT/ATT, ATC/ATG, ACC/GGT, AGC/CTG, AGG/CCT, AAC/GTT, ACT/AGT, CCG/CGG, and ACG/CGT. The 4-nt SSR motifs were mostly ACAT/ATGT, AAAT/ATTT, and AAAG/CTTT, and the 5-nt SSR motifs were represented by AAAAT/ATTTT, AAACC/GGTTT, AAAAG/CTTTT, and AAAAC/GTTTT. The 6-nt SSRs motifs were a minority and, therefore, not shown in Table 1). The output of the MISA tool used to identify the SSRs from the transcriptome and the statistical data are available in “Supplementary file 4 (SSR).txt” and “Supplementary file 5 (MISA).txt” respectively.

**Table S1.** Top 20 simple sequence repeats (SSRs) in a total of 22,549 *D. lanata* SSRs. More than 92% of the total non-redundant SSRs are represented in the above table.

| SSR Motif | No. of repeats | | | | | | | | | | | | Total |
|---|---|---|---|---|---|---|---|---|---|---|---|---|---|
| | 4 | 5 | 6 | 7 | 8 | 9 | 10 | 11 | 12 | 13 | 14 | 15 | >15 | |
| **AG/CT** | - | - | 1503 | 956 | 675 | 453 | 277 | 212 | 142 | 84 | 92 | 73 | 434 | 4901 |
| **AT/AT** | - | - | 1268 | 748 | 626 | 525 | 318 | 527 | 665 | - | - | - | - | 4677 |
| **AC/GT** | - | - | 1093 | 603 | 393 | 293 | 186 | 143 | 99 | 26 | 25 | 20 | 54 | 2935 |
| **AAG/CTT** | - | 776 | 382 | 185 | 80 | 39 | 30 | 28 | 16 | 36 | 22 | 11 | 42 | 1647 |
| **AAT/ATT** | - | 704 | 404 | 171 | 150 | 11 | 35 | 20 | 25 | 13 | 26 | 14 | 27 | 1600 |
| **ATC/ATG** | - | 564 | 225 | 129 | 78 | 28 | 11 | 3 | 2 | 4 | 5 | 5 | 13 | 1067 |
| **ACC/GGT** | - | 505 | 178 | 42 | 23 | 1 | 2 | - | - | - | - | - | 2 | 753 |
| **AGC/CTG** | - | 510 | 193 | 17 | 17 | 1 | - | - | - | - | - | - | - | 738 |
| **AGG/CCT** | - | 366 | 126 | 71 | 28 | 4 | 7 | 9 | 6 | 1 | - | - | - | 618 |
| **AAC/GTT** | - | 179 | 98 | 42 | 25 | 4 | 5 | 5 | 7 | - | - | 2 | 5 | 372 |
| **ACT/AGT** | - | 164 | 83 | 27 | 19 | 15 | 1 | 5 | - | - | - | - | - | 314 |
| **CCG/CGG** | - | 222 | 37 | 19 | - | - | - | - | - | 1 | - | - | - | 279 |
| **ACAT/ATGT** | - | 88 | 39 | 9 | 8 | 16 | 21 | 1 | 3 | 11 | - | - | - | 196 |
| **AAAT/ATTT** | - | 109 | 33 | 2 | 2 | 3 | 4 | 1 | - | - | - | - | - | 154 |
| **AAAAT/ATTTT** | 99 | 18 | 1 | - | 1 | 2 | - | - | - | - | - | - | - | 121 |
| **AAACC/GGTTT** | 82 | 29 | 1 | 4 | 1 | 1 | - | - | - | - | - | - | - | 118 |
| **ACG/CGT** | - | 82 | 14 | 6 | - | - | - | - | - | - | - | - | - | 102 |
| **AAAAG/CTTTT** | 63 | 10 | - | 1 | - | - | - | 5 | - | - | - | - | - | 79 |
| **AAAG/CTTT** | - | 36 | 12 | 5 | - | - | 2 | - | - | - | - | - | - | 55 |
| **AAAAC/GTTTT** | 36 | 9 | 2 | - | - | - | - | - | 1 | - | - | - | - | 48 |
| | | | | | | | | | | | | | Total | 20774 |

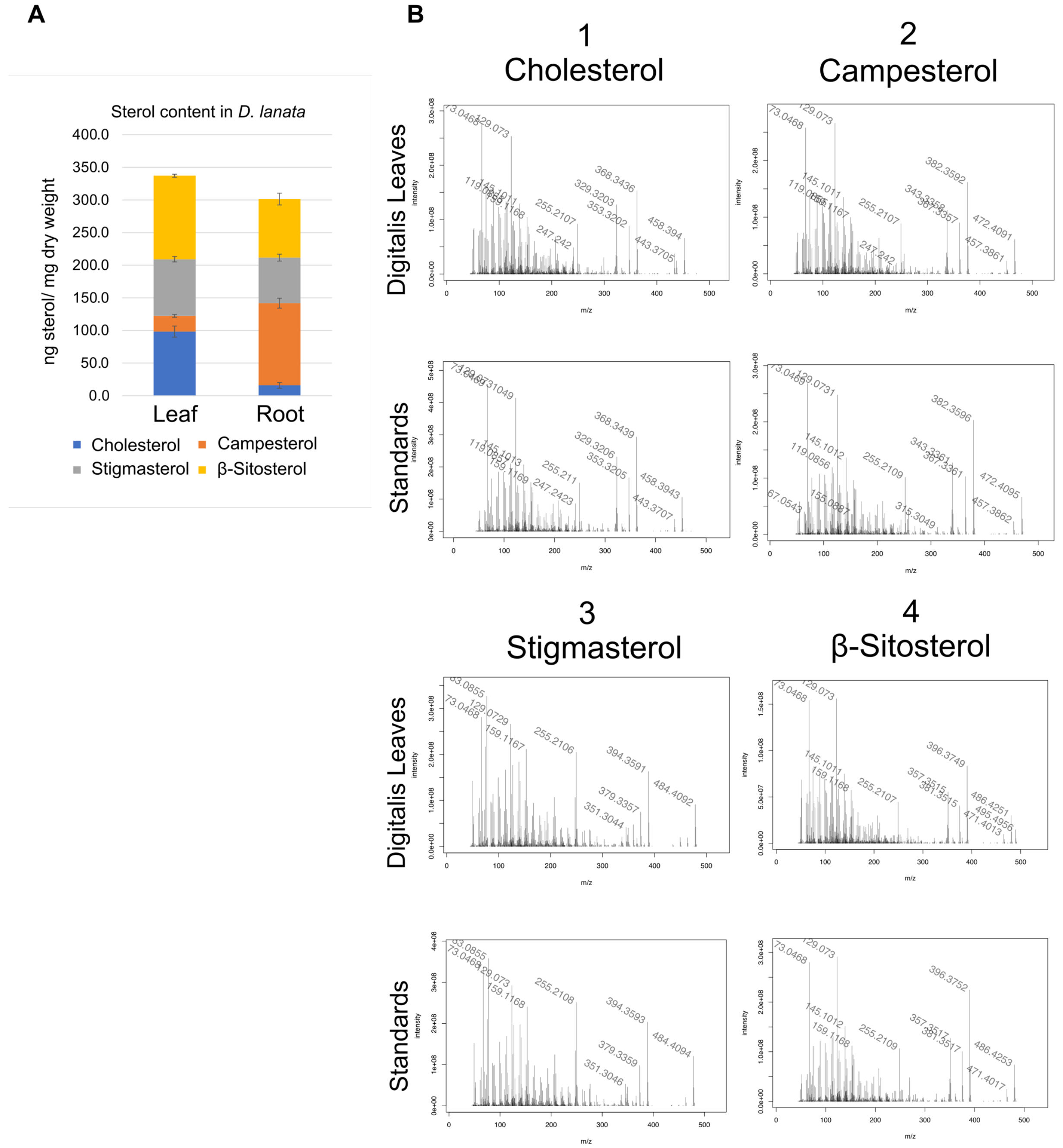


**Figure S5.** GC/MS analysis of steroids in the leaves of *D. lanata*. A) Quantification of sterol contents in *D. lanata* leaves and roots. The sterols are quantified according to Itkin, M., et. al. [33] B) MS spectra of sterols present in *D. lanata* leaves (top) compared to spectra of standards (bottom). Compound 1: cholesterol; compound 2: campesterol; compound 3: stigmasterol; compound 4: *β*-sitosterol.

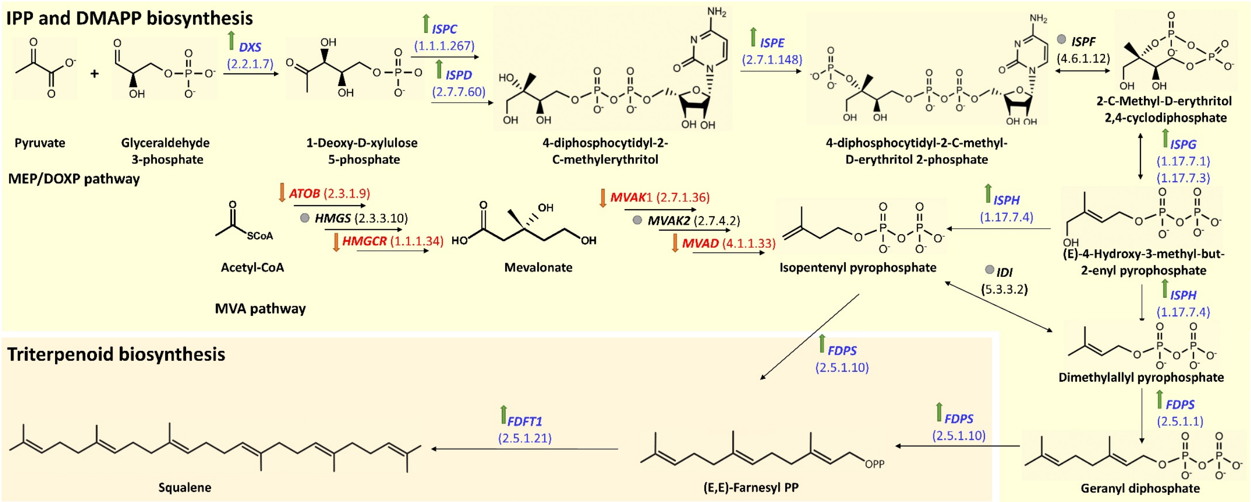


**Figure S6.** Differential expression of terpenoid biosynthetic pathways in leaves vs. roots. The green upward arrow denotes higher expression in leaf versus root; the orange downward arrow conversely denotes higher expression in roots versus leaves; the gray circle denotes almost the same level of expression in roots and leaves. *DXS*: 1-deoxy-D-xylulose-5-phosphate synthase, *ISPC*: 1-deoxy-D-xylulose-5-phosphate reductoisomerase, *ISPD*: 2-C-methyl-D-erythritol 4-phosphate cytidylyltransferase, *ISPE*: 4-diphosphocytidyl-2-C-methyl-D-erythritol kinase, *ISPF* : 2-C-methyl-D-erythritol 2,4-cyclodiphosphate synthase, *ISPG*: (E)-4-hydroxy-3-methylbut-2-enyl-diphosphate synthase, *ISPH* : 4-hydroxy-3-methylbut-2-en-1-yl diphosphate reductase, *HMGS* : hydroxymethylglutaryl-CoA synthase, *MVAK2* : phosphomevalonate kinase, *IDI* : isopentenyl-diphosphate delta-isomerase, *FDPS* : farnesyl diphosphate synthase, *FDFT1* : farnesyl-diphosphate farnesyltransferase.

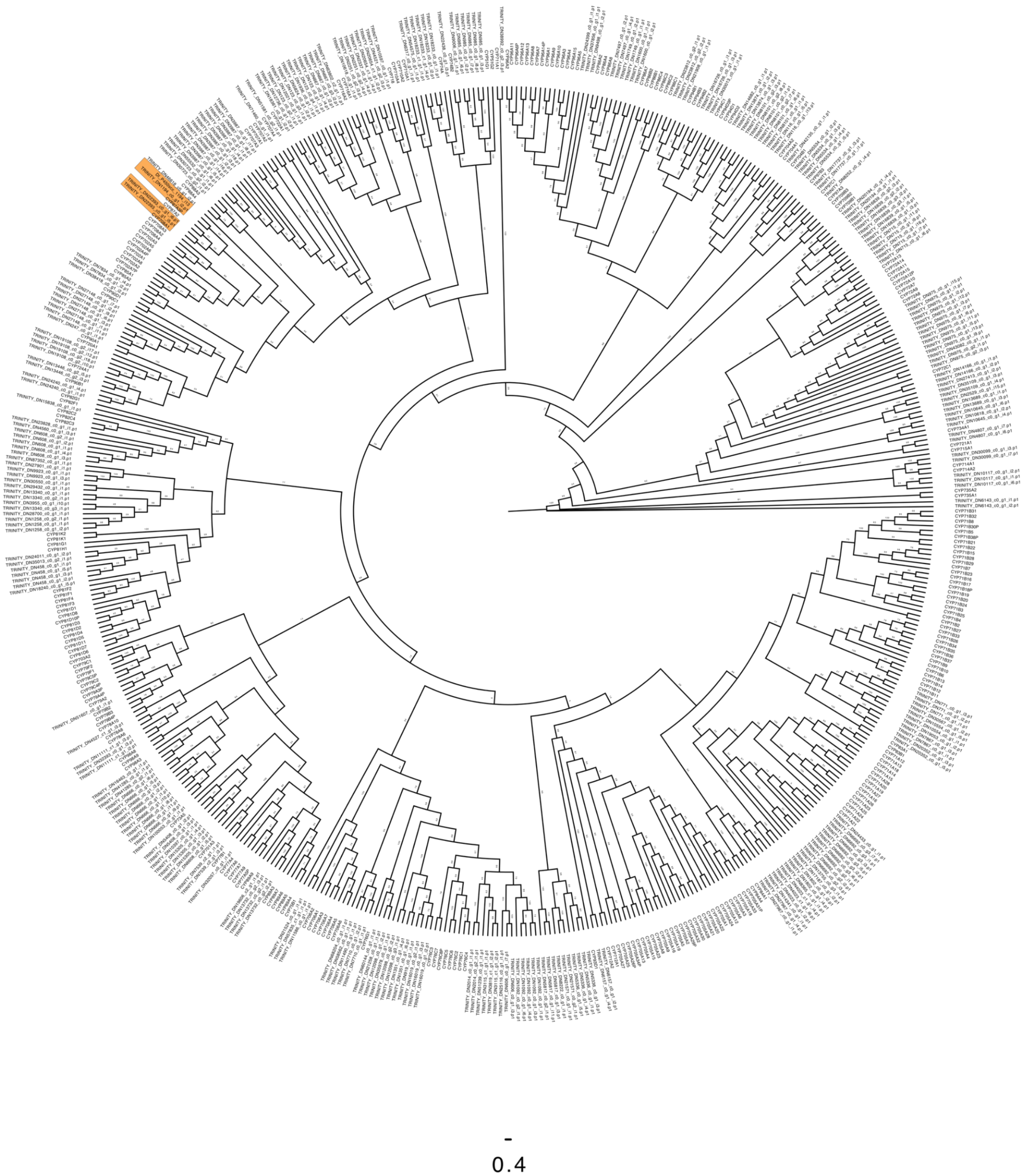


**Figure S7.** Phylogenetic tree with *D. lanata* and *Arabidopsis thaliana* cytochrome P450s. *D. lanata* sequences include transcript that encode proteins containing the Pfam domain PF00067-CYPs and were between 400-600 amino acid long. The Arabidopsis proteins were taken from the *Arabidopsis* cytochrome P450 database [34]. Tree was made with RaxML-NG with proteins aligned in MAFFT.

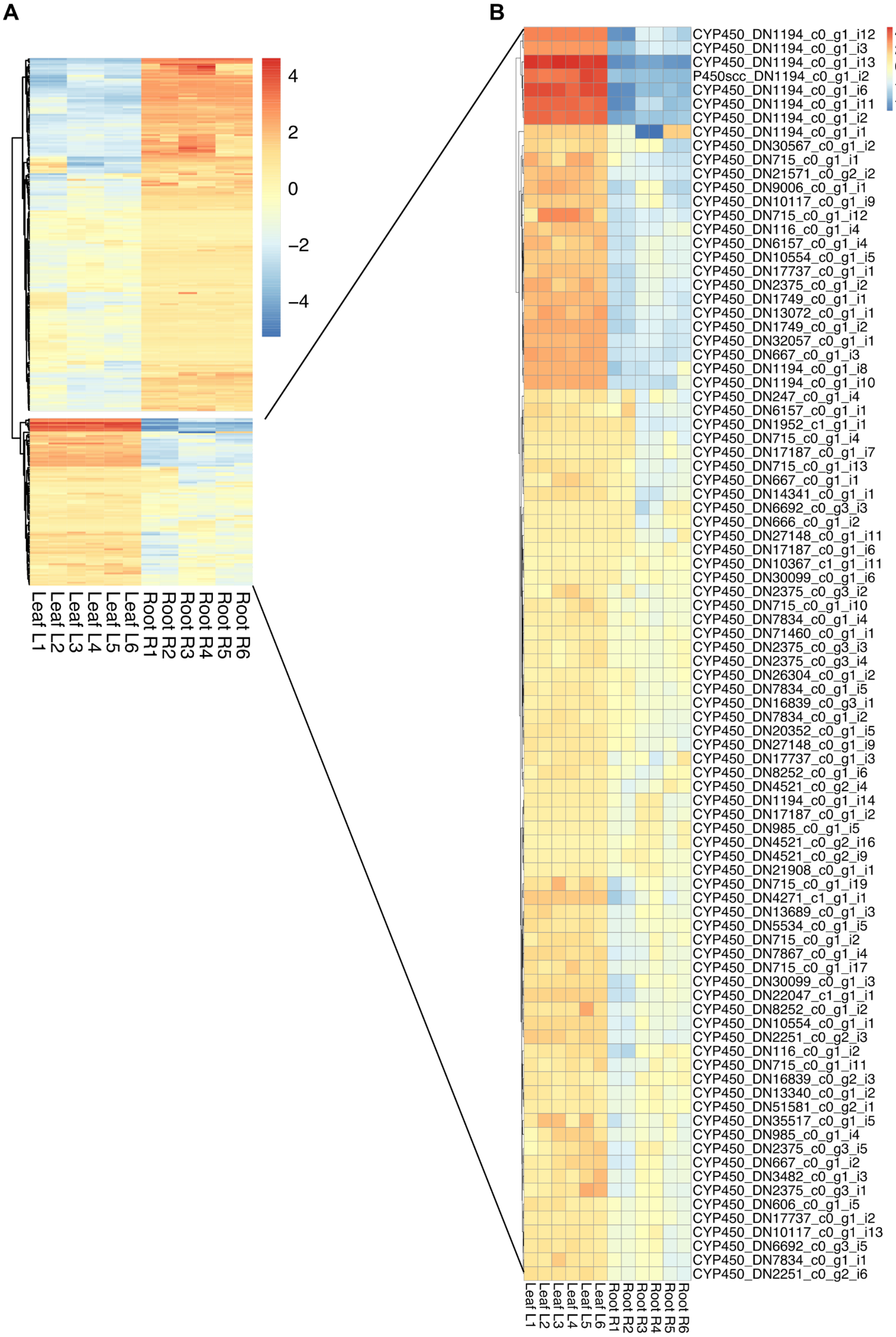


**Figure S8.** Differentially expressed cytochrome P450s in the leaves and roots of *D. lanata*. A) Heatmap of 294 transcripts of cytochrome P450s that are differentially expressed in the leaves and root tissues. Results include six biological and technical replicates. B) 104 cytochrome P450 transcripts are over expressed in leaves than in roots.

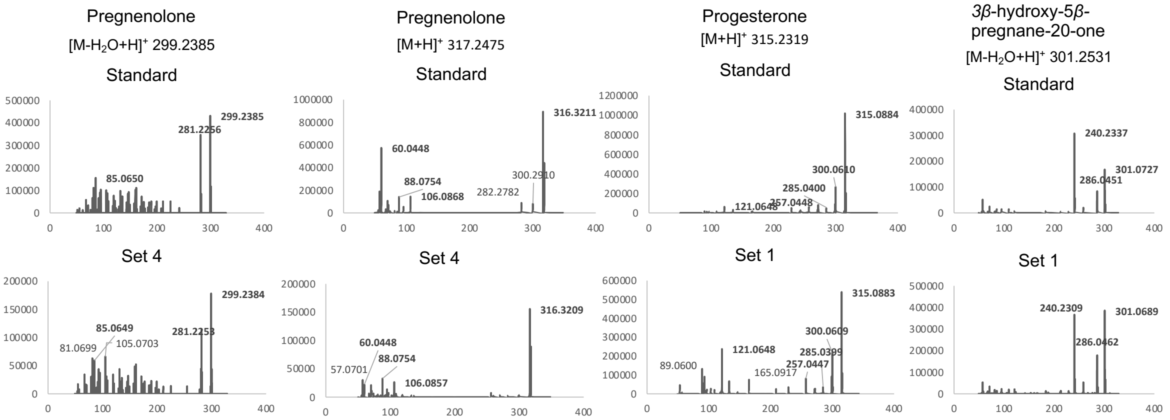


**Figure S9.** MS spectra of pathway intermediates in tobacco transiently expressing digoxin biosynthetic genes in Figure 2C.

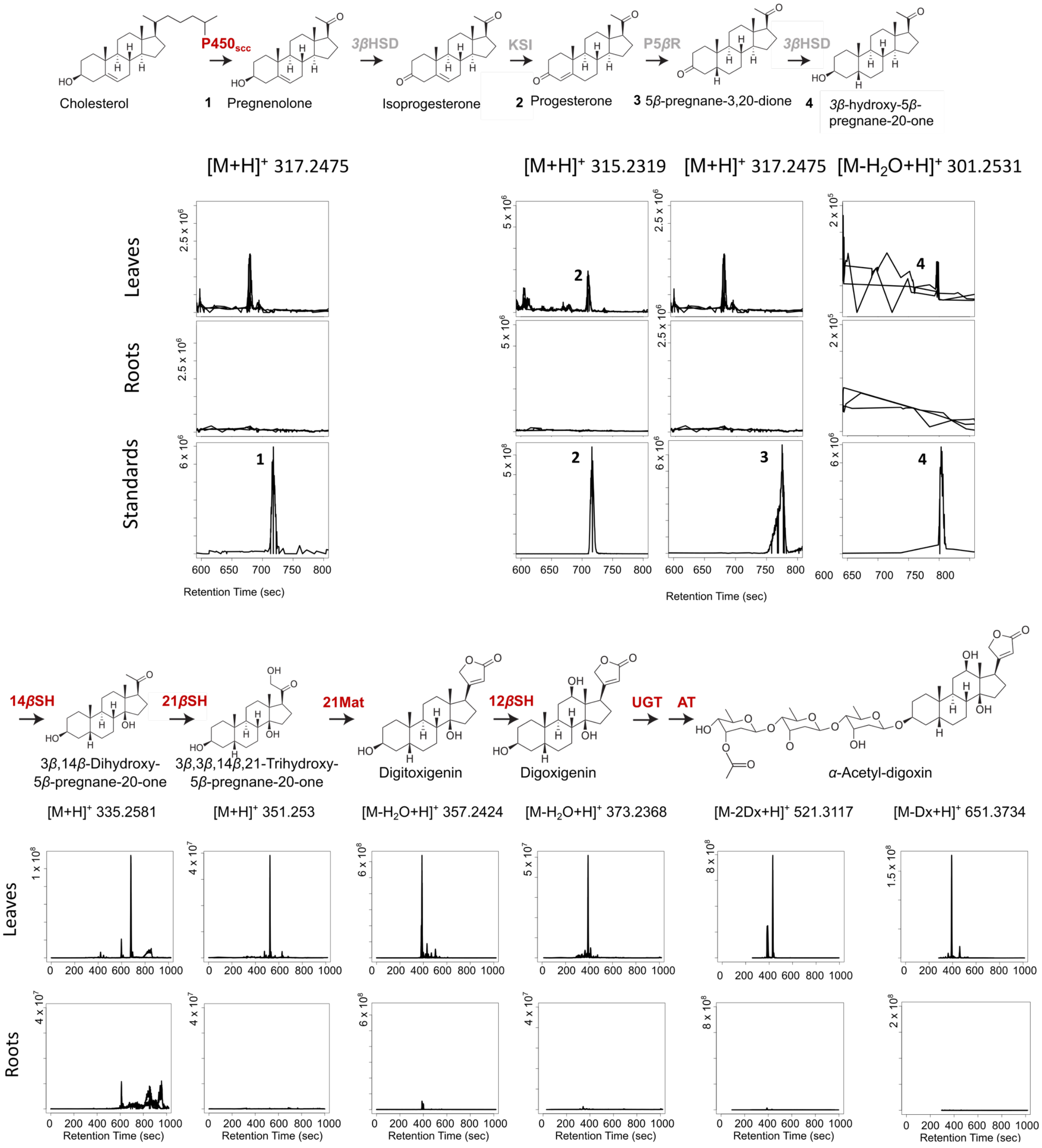


**Figure S10.** Identifying putative digoxin pathway intermediates in *D. lanata*. Extracted ion chromatograms for each compound in the pathway. Standards are provided for pregnenolone (1), progesterone (2), 5*β*-pregnane-3,20-one (3), and 3*β*-hydroxy-5*β*-pregnane-20-one (4). Dx: digitoxose unit.

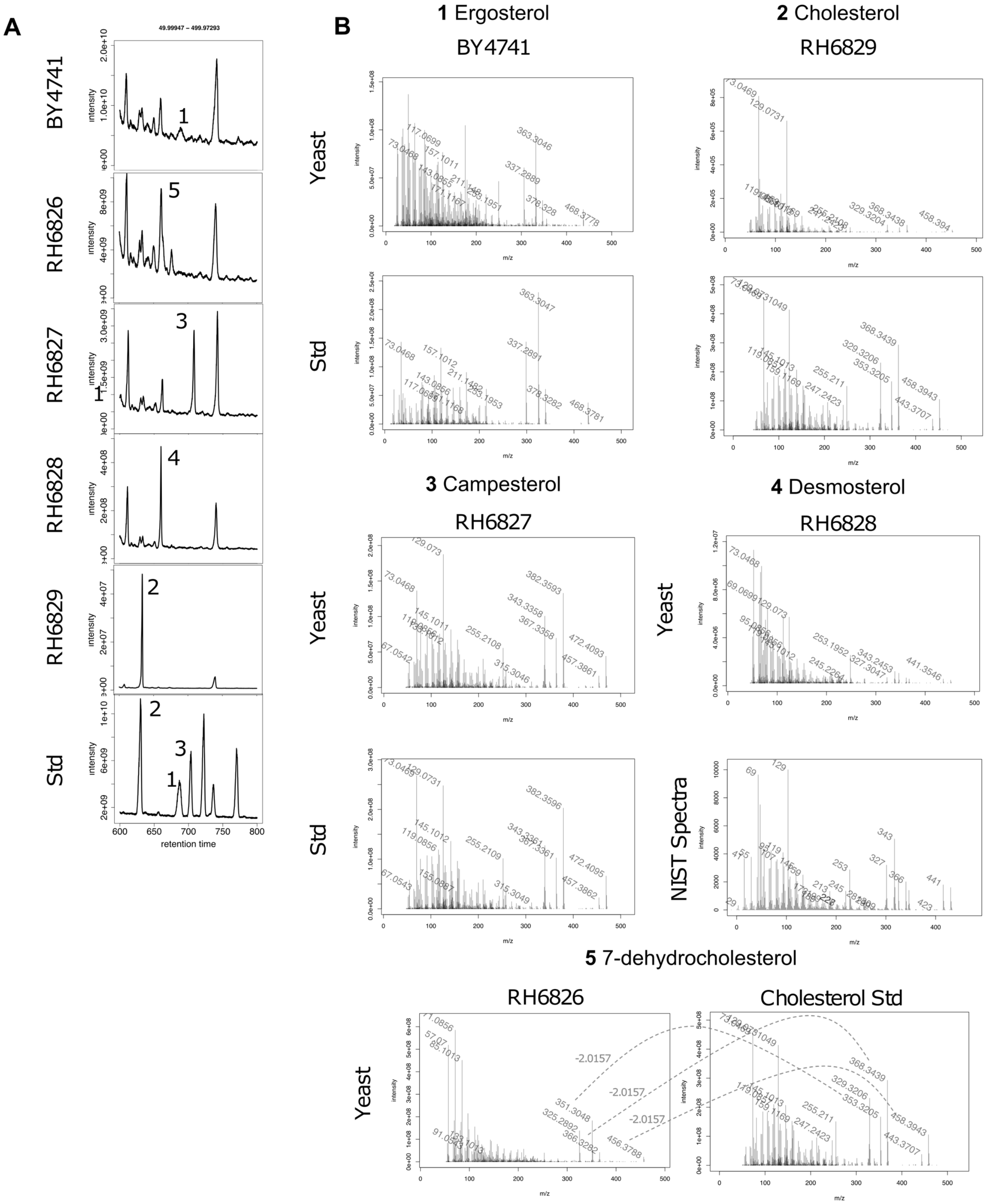


**Figure S11.** GC/MS analysis of sterol contents in engineered yeast strains. BY4741: wildtype yeast containing ergosterol; RH6826: 7-dehydrocholesterol producing yeast, RH6827: campesterol-producing yeast; RH6828: desmosterol-producing yeast; RH6829: cholesterol-producing yeast. A) Total ion chromatogram of yeast strains compared to authentic standards. Compound 1: ergosterol; compound 2: cholesterol, compound 3: campesterol; compound 4: desmosterol; compound 5: 7-dehydrocholesterol. B) MS spectra of indicated peaks compared to authentic standards or spectra in NIST database.

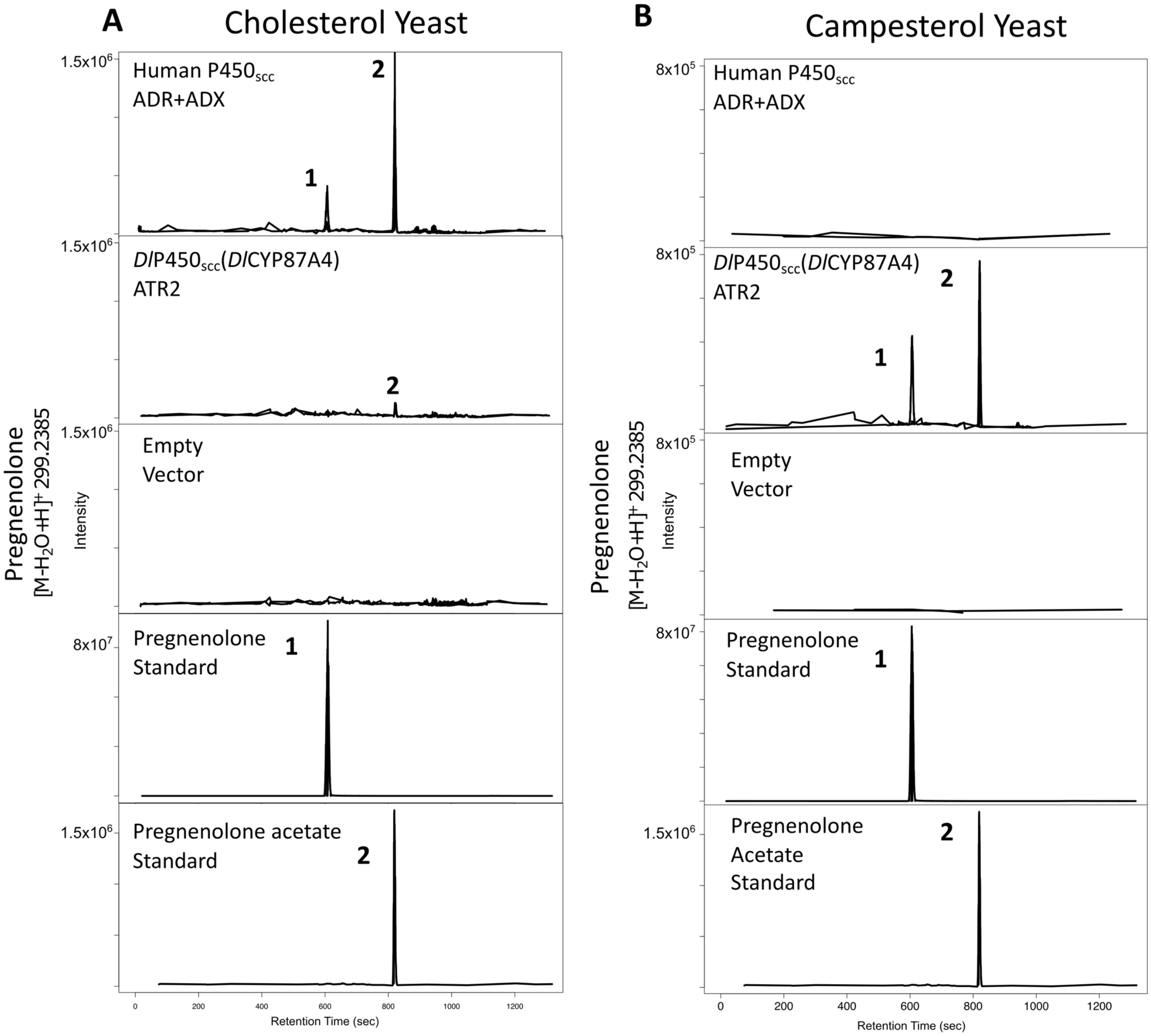


**Figure S12.** Preliminary experiment with campesterol-producing yeast expressing *Dl*CYP87A4. A) Cholesterol producing yeast strain (RH6829) expressing either the human $P450_{scc}$ or the *Digitalis* $P450_{scc}$ with their respective redox partners. Yeast was harvested after growing in shake flask for 21 hours followed by extraction and LC/MS. ADX: adrenodoxin, ADR: adrenodoxin reductase. ATR2: *Arabidopsis* cytochrome P450 reductase 2. B) Campesterol-producing yeast strain (RH6827) expressing either the human $P450_{scc}$ or the *Digitalis* $P450_{scc}$ with their respective redox partners. Yeast was harvested after growing in shake flask for 18 hours followed by extraction and LC/MS. Compound 1: pregnenolone; compound 2: pregnenolone acetate.

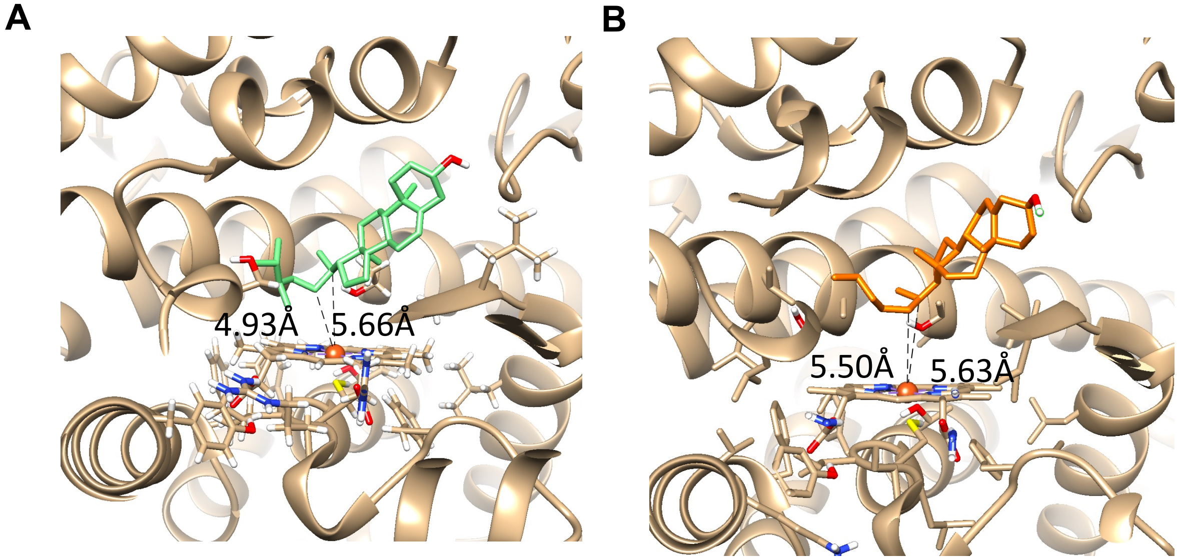


**Figure S13.** Docking of campesterol (A) and cholesterol (B) into the active site of *Dl*CYP87A4 model. The dashed lines measure the distances between the C20 or C22 carbons and the iron center in the heme. The heme is supposed to add an oxygen atom to these two carbons.

**Table S2.** *Dl*CYP87A4 with closely related cytochrome P450s in various plant species. Aa: amino acid.

| Query | Subject | Species | % Identical (aa) | Alignment Length | Mismatches | Gap Opens | E Value | Bit Score |
|---|---|---|---|---|---|---|---|---|
| DlCYP87A4 | dpa_locus_4690_iso_4_len_1619_ver_1 | *Digitalis purpurea* | 97.30 | 408 | 10 | 1 | 0 | 827 |
| DlCYP87A4 | XP_011081025.1 | *Sesamum indicum* | 74.42 | 473 | 116 | 3 | 0 | 747 |
| DlCYP87A4 | XP_022881263.1 | *Olea europaea* | 74.63 | 469 | 114 | 3 | 0 | 744 |
| DlCYP87A4 | *Dl*CYP87A1 | *Digitalis lanata* | 72.27 | 476 | 129 | 2 | 0 | 727 |
| DlCYP87A4 | cal_g005561.t1 | *Calotropis gigantea* | 68.99 | 445 | 133 | 3 | 0 | 636 |
| DlCYP87A4 | m_44857 | *Asclepias syriaca* | 52.49 | 482 | 214 | 6 | 0 | 530 |
| DlCYP87A4 | m_12618 | *Asclepias syriaca* | 38.57 | 446 | 263 | 7 | 2.38E-111 | 330 |
| DlCYP87A4 | *Dl*CYP87A3 | *Digitalis lanata* | 97.45 | 470 | 11 | 1 | 0 | 891 |
| DlCYP87A4 | *Dl*CYP87A2 | *Digitalis lanata* | 74.18 | 457 | 115 | 2 | 0 | 692 |

**Table S3.** Primers used in this study. FWD/FW/F: forward primers, REV/RW/R: reverse primers. Capital letters: gene-specific nucleotides; Small case letters: additional nucleotides for cloning.

| Name | Sequence |
|---|---|
| **Primers for cloning genes into the pEAQ vector by Gibson Cloning** | |
| TRINITY_DN247_c0_g1_i1.p1 FWD | tattctgcccaaattcgcgaATGGACTCCAACATGTTTCTCTAC |
| TRINITY_DN247_c0_g1_i1.p1 REV | accagagttaaaggcctcgaTCACTTAAGCCTCTGAATGATAATCG |
| TRINITY_DN1194_c0_g1_i13.p1 FWD | tattctgcccaaattcgcgaATGTCGTTAGTAGCTATGAGC |
| TRINITY_DN1194_c0_g1_i13.p1 REV | accagagttaaaggcctcgaTTAGCCTCTCTCAGTCATGT |
| 3βHSD_FWD | tattctgcccaaattcgcgaATGTCGTCAAAGCCAAGGTTGG |
| 3βHSD_REV | accagagttaaaggcctcgaCTAACGCACGACGGTGAAGC |
| P5βR-2_FWD | tattctgcccaaattcgcgaATGTATACCGACACAACGACT |
| P5βR-2_REV | accagagttaaaggcctcgaTCAAGGGACAAATCTATAAGATCTCA |
| **Primers for Golden Gate cloning and domestication into the pYTK001 vector** | |
| 1194 + pYTK001 F | tttcgtctcatcggggtctcatATGTCGTTAGTAGCTATGAGC |
| 1194 sub 1 R | TCCCGTCTCTCATGTAGCATA |
| 1194 sub 2 F | tttcgtctctCATGAGAGgCGGGAAAAGC |
| 1194 sub 2 R | tttcgtctcAGTGAGAGCTTTT |
| 1194 sub 3 F | tttcgtctctTCACTGAaACCGAATTTAAAGG |
| 1194 +pYTK001 R | tttcgtctcaggtcggtctcaggaTTAGCCTCTCTCAGTCATGT |
| **Primers for qRTPCR** | |
| TRINITY_DN1194_c0_g1_i13.p1 1110-1131 FWD 1 | ATACACAATTCCAGCTGGTTGG |
| TRINITY_DN1194_c0_g1_i13.p1 1285-1306 REV 1 | TCTGGACCTTTGTGAAATCTGC |
| DN247 qRT FW | CCACACGCATTGGCTCATCTTAAG |
| DN247 qRT RW | CGACACATTGAGTGAACGGCATAG |

**Table S4.** Yeast strains used in this study. The RH strains are from Dr. Howard Riezman's lab [35].

| Strain | Genotype |
|---|---|
| RH6826 | MATa *ura3 leu2 his3 trp1 can1 bar1 erg5Δ::HIS5-TDH3-DHCR24 erg6Δ::TRP1* |
| RH6827 | MATa *ura3 leu2 his3 trp1 can1 bar1 erg5Δ::TRP1-TDH3-DHCR7* |
| RH6828 | MATα *ura3 lue2 trp1 ade2 can1 bar1 erg6Δ::TRP1-TDH3-DHCR7* |
| RH6829 | MATa *ura3 leu2 his3 trp1 can1 bar1 erg5Δ::HIS5-TDH3-DHCR24 erg6Δ::TRP1-TDH3-DHCR7* |
| BY4741 | MATa *his3Δ1 leu2Δ0 met15Δ0 ura3Δ0* |

**Table S5**. Constructs used in this study. The DNA parts for transcription-unit-level and multi-gene-level plasmids care described in Lee, et.al (2015) [36].

| Plasmid | Description |
|---|---|
| pEAQ-HT_*3βHSD* | pEAQ vector cloned with *D. lanata* gene "3βHSD". Transcript ID: "TRINITY_DN638_c0_g3_i1.p1" |
| pEAQ-HT_*Pβ5R-2* | pEAQ vector cloned with *D. lanata* gene "P5βR-2". NCBI accession: ADL28122.1 |
| pEAQ-HT_*DlCYP90A1-P450* | pEAQ vector cloned with *D. lanata* gene "P450". Transcript ID: "TRINITY_DN247_c0_g1_i1.p1" |
| pEAQ-HT_*CYP-DlCYP87A4* | pEAQ vector cloned with *D. lanata* putative gene as a cytochrome P450 steroid hydroxylase or $P450_{scc}$. |
| pYTK001_*HsP450scc* | Human $P450_{scc}$, or CYP11A1, codon-optimized for yeast expression |
| pYTK001_*HsADX* | Human andrenodoxin (ADX), codon-optimized for yeast expression |
| pYTK001_*HsADR* | Human andrenodoxin reductase (ADX), codon-optimized for yeast expression |
| pYTK001_*DlCYP87A4* | *Digitalis lanata* $P450_{scc}$ TRINITY_DN1194_c0_g1_i13 domesticated for cloning into the pYTK001 |
| pYTK001_*ATR2* | *Arabidopsis thaliana* cytochrome P450 reductase 2 (ATR2) cloned into pYTK001 |
| pTU11S1_*DlCYP87A4* | Transcription-unit-level plasmid of *D. lanata* DN1194 with the pTDH3 promoter, the tENO1 terminator, and LS-R1 connectors |
| pTU1116_ATR2 | Transcription-unit-level plasmid of ATR2 with the pCCW12 promoter, the tSSA1 terminator, and L1-RE connectors |
| pTU11S1_*HsP450scc* | Transcription unit with promoter-CDS-terminator for Human P450scc and LE-R1 linker |
| pTU1112_*HsADX* | Transcription unit with promoter-CDS-terminator for human ADX and L1-R2 linker |
| pTU1126_*HsADR* | Transcription unit with promoter-CDS-terminator for human ADR and L2-RE linker |
| pMGR22_*DlCYP87A4.ATR2* | Multigene plasmid with *D. lanata* TRINITY_DN1194_c0_g1_i13 + ATR2 |
| pMGR22_*HsP450scc.HsADX.ADR* | Multigene plasmid with Human $CPY450_{scc}$ (CYP11A1) and Human redox partners ADR and ADX |

***References***

1 Andrews, S. *FastQC: a quality control tool for high throughput sequence data*, <http://www.bioinformatics.babraham.ac.uk/projects/fastqc> (2010).

2 Haas, B. J. *et al.* De novo transcript sequence reconstruction from RNA-seq using the Trinity platform for reference generation and analysis. *Nature protocols* **8**, 1494 (2013).

3 Langmead, B. & Salzberg, S. L. Fast gapped-read alignment with Bowtie 2. *Nature methods* **9**, 357-359 (2012).

4 Li, W. & Godzik, A. Cd-hit: a fast program for clustering and comparing large sets of protein or nucleotide sequences. *Bioinformatics* **22**, 1658-1659 (2006).

5 Altschul, S. F., Gish, W., Miller, W., Myers, E. W. & Lipman, D. J. Basic local alignment search tool. *Journal of molecular biology* **215**, 403-410 (1990).

6 Pruitt, K. D., Tatusova, T. & Maglott, D. R. NCBI reference sequences (RefSeq): a curated non-redundant sequence database of genomes, transcripts and proteins. *Nucleic acids research* **35**, D61-D65 (2007).

7 Consortium, U. The universal protein resource (UniProt) in 2010. *Nucleic acids research* **38**, D142-D148 (2010).

8 Finn, R. D. *et al.* Pfam: the protein families database. *Nucleic acids research* **42**, D222-D230 (2014).

9 Quevillon, E. *et al.* InterProScan: protein domains identifier. *Nucleic acids research* **33**, W116-W120 (2005).

10 Ye, J. *et al.* WEGO 2.0: a web tool for analyzing and plotting GO annotations, 2018 update. *Nucleic acids research* **46**, W71-W75 (2018).

11 Li, B. & Dewey, C. N. RSEM: accurate transcript quantification from RNA-Seq data with or without a reference genome. *BMC bioinformatics* **12**, 323 (2011).

12 Robinson, M. D. & Oshlack, A. A scaling normalization method for differential expression analysis of RNA-seq data. *Genome biology* **11**, R25 (2010).

13 Wagner, G. P., Kin, K. & Lynch, V. J. Measurement of mRNA abundance using RNA-seq data: RPKM measure is inconsistent among samples. *Theory in biosciences* **131**, 281-285 (2012).

14 Pérez-Rodríguez, P. *et al.* PlnTFDB: updated content and new features of the plant transcription factor database. *Nucleic acids research* **38**, D822-D827 (2010).

15 Jin, J., Zhang, H., Kong, L., Gao, G. & Luo, J. PlantTFDB 3.0: a portal for the functional and evolutionary study of plant transcription factors. *Nucleic acids research* **42**, D1182-D1187 (2014).

16 Lehti-Shiu, M. D. & Shiu, S.-H. Diversity, classification and function of the plant protein kinase superfamily. *Philosophical Transactions of the Royal Society B: Biological Sciences* **367**, 2619-2639 (2012).

17 Thiel, T., Michalek, W., Varshney, R. & Graner, A. Exploiting EST databases for the development and characterization of gene-derived SSR-markers in barley (Hordeum vulgare L.). *Theoretical and applied genetics* **106**, 411-422 (2003).

18 Manni, M., Berkeley, M. R., Seppey, M., Simão, F. A. & Zdobnov, E. M. BUSCO update: novel and streamlined workflows along with broader and deeper phylogenetic coverage for scoring of eukaryotic, prokaryotic, and viral genomes. *Molecular Biology and Evolution* **38**, 4647-4654 (2021).

19 MacManes, M. D. On the optimal trimming of high-throughput mRNA sequence data. *Frontiers in genetics* **5**, 13 (2014).

20 Yang, L. *et al.* Transcriptome analysis of medicinal plant Salvia miltiorrhiza and identification of genes related to tanshinone biosynthesis. *PLoS One* **8** (11), e80464 (2013).

21 Gupta, P. *et al.* De novo assembly, functional annotation and comparative analysis of Withania somnifera leaf and root transcriptomes to identify putative genes involved in the withanolides biosynthesis. *PLoS One* **8** (5), e62714 (2013).

22 Wu, Y.-H. *et al.* Comparative analysis of the chrysanthemum leaf transcript profiling in response to salt stress. *PloS one* **11** (7), e0159721 (2016).

23 Pandey, A. *et al.* Transcriptome and metabolite analysis reveal candidate genes of the cardiac glycoside biosynthetic pathway from Calotropis procera. *Scientific reports* **6**, 34464 (2016).

24 Chabikwa, T. G., Barbier, F. F., Tanurdzic, M. & Beveridge, C. A. De novo transcriptome assembly and annotation for gene discovery in avocado, macadamia and mango. *Scientific Data* **7** , 9 (2020).

25 Langmead, B., Trapnell, C., Pop, M. & Salzberg, S. L. Ultrafast and memory-efficient alignment of short DNA sequences to the human genome. *Genome biology* **10**, R25 (2009).

26 Wang, Z., Gerstein, M. & Snyder, M. RNA-Seq: a revolutionary tool for transcriptomics. *Nature reviews genetics* **10**, 57-63 (2009).

27 Kanehisa, M. & Sato, Y. KEGG Mapper for inferring cellular functions from protein sequences. *Protein Science* **29**(1), 28-35 (2019).

28 Gish, L. A. & Clark, S. E. The RLK/Pelle family of kinases. *The Plant Journal* **66**, 117-127 (2011).
29 Eisen, J. A., Sweder, K. S. & Hanawalt, P. C. Evolution of the SNF2 family of proteins: subfamilies with distinct sequences and functions. *Nucleic acids research* **23**, 2715-2723 (1995).
30 Sun, X., Wang, Y. & Sui, N. Transcriptional regulation of bHLH during plant response to stress. *Biochemical and biophysical research communications* **503**, 397-401 (2018).
31 Chen, H. *et al.* Transcriptome sequencing of mung bean (Vigna radiate L.) genes and the identification of EST-SSR markers. *PloS one* **10** (4), e0120273 (2015).
32 Gupta, P. K. *et al.* Transferable EST-SSR markers for the study of polymorphism and genetic diversity in bread wheat. *Molecular genetics and genomics* **270**, 315-323 (2003).
33 Itkin, M. *et al.* GLYCOALKALOID METABOLISM1 is required for steroidal alkaloid glycosylation and prevention of phytotoxicity in tomato. *The Plant Cell* **23**, 4507-4525 (2011).
34 Paquette, S. M., Bak, S. & Feyereisen, R. Intron–exon organization and phylogeny in a large superfamily, the paralogous cytochrome P450 genes of Arabidopsis thaliana. *DNA and cell biology* **19**, 307-317 (2000).
35 Souza, C. M. *et al.* A stable yeast strain efficiently producing cholesterol instead of ergosterol is functional for tryptophan uptake, but not weak organic acid resistance. *Metabolic engineering* **13**, 555-569 (2011).
36 Lee, M. E., DeLoache, W. C., Cervantes, B. & Dueber, J. E. A highly characterized yeast toolkit for modular, multipart assembly. *ACS synthetic biology* **4**, 975-986 (2015).
